# ENOD93 interacts with cytochrome c oxidase altering respiratory ATP production and root growth in plants

**DOI:** 10.1101/2023.04.15.535782

**Authors:** Chun Pong Lee, Xuyen H. Le, Ryan M.R. Gawryluk, José A. Casaretto, Steven J. Rothstein, A. Harvey Millar

## Abstract

The early nodulin 93 (ENOD93) gene family in plants can regulate biological nitrogen fixation in legumes and nitrogen use efficiency in cereals but its molecular function is unknown. We show profile hidden Markov models define ENOD93 as a distant homolog of the N-terminal domain of RESPIRATORY SUPERCOMPLEX FACTOR 2 (RCF2). RCF2 is reported to regulate cytochrome oxidase (CIV) influencing the generation of a mitochondria proton motive force in yeast. Knockout of *enod93* in Arabidopsis leads to a short root phenotype. ENOD93 is associated with a protein complex the size of CIV in isolated mitochondria but neither CIV abundance nor its activity in ruptured organelles changed in *enod93*. However, a progressive loss of ADP-dependent respiration rate was observed in *enod93* mitochondria which could be fully recovered in complemented lines. Mitochondrial membrane potential was higher in *enod93* but ATP synthesis and ADP depletion rates progressively decreased. Respiration rate of whole *enod93* seedlings was elevated and root ADP content was nearly double that in WT without a change in ATP content. These altered energetic states correlated with elevated respiratory substrate levels in roots of *enod93* compared to WT and complemented lines. Overexpression of ENOD93 lowered ATP content in roots and increased the abundance of a range of amino acids in both roots and leaves. We propose that two previously unconnected gene families in plants, ENOD93 and HYPOXIA INDUCED GENE DOMAIN, are the functional equivalent of yeast RCF2 but have remained undiscovered in many eukaryotic lineages because they are encoded in two separate genes.

**Highlight significance:** We identify the enigmatic early nodulin ENOD93 gene family as the plant homolog of the N-terminal regulatory domain of the yeast RESPIRATORY SUPERCOMPLEX 2 (RCF2) of the mitochondrial oxidative phosphorylation system and provide biochemical and physiological evidence of its role in plant ATP production, broadly explaining the role of ENOD93 in plants.

## Introduction

Early nodulins (ENOD) were first discovered 30 years ago as a diverse group of genes expressed at early stages of nodule development in pea, alfalfa and soybean (Kouchi and Hata, 1993; Pichon et al., 1992; Scheres et al., 1990; Yang et al., 1993). ENOD homologs have subsequently been found in non-leguminous plants (Oldroyd and Downie, 2008) and diverse roles in plant biology as membrane proteins promoting the exchange of sugars, amino acids, cofactors and nutrients have been proposed, with differing levels of supporting evidence in each case (Denance et al., 2014). One of the most enigmatic of the ENOD set of genes has been ENOD93 for which a molecular function remains unknown. It was first identified in soybean as a gene encoding a 105 amino acid hydrophobic protein rich in alanine and serine residues (Kouchi and Hata, 1993). Expression of soybean *ENOD93* during nodulation has subsequently been shown to be controlled by a specific miRNA and loss of ENOD93 expression is a control point in soybean nodule formation and biological N_2_ fixation (Yan et al., 2015). However, ENOD93-like proteins are not restricted to legumes but are present in nearly all plant genomes and are often present in small gene families. Reddy et al. (Reddy et al., 1999; Reddy et al., 1998) identified a series of ENOD93 homologs in rice. From the rice gene family of 7 members, OsENOD93-1 was prioritized for investigation as a nitrogen use efficiency (NUE) gene candidate because of its strong transcriptional response to both decreasing and increasing N-conditions, suggesting it was a central regulator responding to N fluctuation (Bi et al., 2007). Transgenic plants over-expressing OsENOD93-1 accumulated amino acids in roots and in xylem sap better than WT under low and medium N-conditions and show increased shoot dry biomass and seed yield compared to WT plants under moderate N-conditions (Bi et al., 2009).

ENOD93 is the only member of the ENOD gene set predicted to be located inside mitochondria in plants based on sequence prediction algorithms and subcellular location experiments (Hooper et al., 2014). Expression of a C-terminal fusion of OsENOD93-1 to YFP experimentally confirmed it was located inside mitochondria (Bi et al., 2009). In Arabidopsis, a single ENOD93 homolog is found in the nuclear genome encoded by the gene At5g25940. Like its rice counterpart, it is also predicted to be located inside mitochondria (Hooper et al., 2017). AtENOD93 has been found in the Arabidopsis mitochondrial proteome by peptide mass spectrometry in five independent literature reports (Brugiere et al., 2004; Heazlewood et al., 2004; Klodmann et al., 2011; Senkler et al., 2017; Taylor et al., 2011). While ENOD93 is only a small 12 kDa protein, it has been observed in native gels at a molecular mass of hundreds of kilodaltons, suggesting ENOD93 is part of a larger protein complex (Klodmann et al., 2011; Senkler et al., 2017).

As the function of ENOD93 is unknown and has eluded determination for decades, we sought to further investigate its sequence similarity, localisation within the mitochondria and molecular function. In so doing we aimed to link the genetic evidence for its role in nitrogen-fixing nodules and in nitrogen use efficiency to its presence and role as a conserved mitochondrial membrane protein in plants. We conclude that ENOD93 is a divergent supernumerary subunit of cytochrome c oxidase (CIV). It is related to the N-terminal domain of RESPIRATORY SUPERCOMPLEX FACTOR 2 in yeast and it regulates the ability of CIV in plants to operate under a protonmotive force to enable efficient oxidative phosphorylation in mitochondria. The high demand for mitochondrial ATP during nitrogen-linked processes likely explains its previous links to these processes in plants.

## Results

### ENOD93 sequence conservation among eukaryotes and similarity to the N-terminus of yeast RCF2

ENOD93 sequences have been widely found in sequenced plant genomes and conservation of their core domain is enshrined in the ENOD93 Profam domain (PF03386) **(Figure S1A)**. Proteins with regions similar to ENOD93 fall into two general physicochemical classes. The first class, found in plants, some algae, and some microbial eukaryotes (*e.g.*, amoebozoans), was made up of proteins approximately 90-130 amino acids in length, typically with two predicted transmembrane helices or areas of increased hydrophobicity **(Figure S2)**. Proteins in the other class, found in stramenopiles, haptophytes and basal holozoans, were typically longer than 200 amino acids, with four predicted transmembrane helices or regions of increased hydrophobicity. In the latter case, homologs generally had an N-terminal region similar to ENOD93 and an additional C-terminal portion containing a conserved “Hig_1_N” domain thought to be involved in the cellular response to hypoxia. The opposite orientation of regions similar to ENOD93 and Hig_1_N was noted in provorans (Tikhonenkov et al., 2022) and bolidophytes.

Blast-based inspection of putative holozoan ENOD93 homologs including the conserved Hig_1_N domain revealed similarity to yeast RCF2 (respiratory super<u>c</u>omplex factor family 2), a protein that plays a role in the regulation of cytochrome c oxidase (Complex IV) in the inner mitochondrial membrane (Strogolova et al., 2012). As found in other holozoans and protists with joined ENOD93- like and Hig_1_N regions, yeast RCF2 is composed of a C-terminal portion with two transmembrane helices and a Hig_1_N domain, and an N-terminal region lacking obvious similarity to other known protein domains, but that profile HMM searches show to be homologous to ENOD93 (**Figure S2**).

Given that plant-like ENOD93 homologs are similar only to the N-terminal portion of fungal RCF2, we hypothesized that plant ENOD93s represent separate genes analogous to the N-terminal region of fungal RCF2, and that plants – and microbial eukaryotes with plant-like ENOD93 homologs – likely encode separate and shorter RCF2 proteins that include the Hig_1_N domain. To this end, we identified putative ‘truncated’ Hig_1_N domain-containing RCF2 homologs that are similar to the C-terminal region of fungal RCF2 across plants and in several microbial eukaryotes **(Figures S1B, S2B**). The *Arabidopsis* RCF2 candidates, ATHIGD2 (At5g27760) and ATHIGD3 (At3g05550), were already known to possess Hig_1_N domains and we had previously proposed them as plant-specific components of cytochrome *c* oxidase (Millar et al., 2004). However, they were not recognised at that time to be potential homologs of the C-terminus of RCF2.

In yeast it is known that RCF2 is proteolytically cleaved *in vivo* into separate N- and C-terminal fragments (Rompler et al., 2016) that are roughly equivalent to the ENOD93-like and C-terminal Hig_1_N regions, suggesting that these proteins may act as physically separate polypeptides even when encoded in a single open reading frame. Yeast encodes an additional mitochondrial protein bearing a Hig_1_N domain, RCF1, which functions in maturation of cytochrome *c* oxidase (Strogolova et al., 2012). RCF1 has an *Arabidopsis* homolog (Hwang et al., 2017), ATHIGD1 (AT3G48030.1), and is well conserved in terms of distribution and sequence across eukaryotes, including animals. Yeast also encodes another protein, RCF3, that is homologous to the N-terminal region of RCF2 (Rompler et al., 2016). However, plant ENOD93 is more similar to the N-terminal region of yeast RCF2 than to RCF3, and RCF3 homologs could not be identified with confidence outside of Ascomycota and Mucoromycota. This suggests that RCF3 is the product of a more recent gene duplication event within Fungi.

Pairwise profile HMM comparisons show the relative similarity (E-value) between Arabidopsis HIG1, HIG2 and ENOD93, yeast RCF1 and RCF2, and human HIGD1A/2A sequences **(Figure 1A)**. Collectively, these phylogenetic data suggest the existence of separate RCF1 like proteins in plants, yeast and mammals which contain a Hig_1_N domain, while homologs of the RCF2 like protein in yeast exist as two separate genes in plants and algae **(Figure 1B)**. While the homology of ENOD93 proteins is readily apparent between plants (Pfam PF03386), there are few sequence features conserved broadly amongst eukaryotes. Nonetheless, there are several highly conserved amino acids, which likely occur near the start or end of transmembrane helices defined for the N-terminal region of yeast RCF2 (Zhou et al., 2021). Sequence similarity of standalone plant-type RCF2 and the fungal-type RCF2 C-terminal region is more apparent, with multiple highly conserved residues at the borders of, and within regions that are transmembrane helices in yeast **(Figure 1C, Figure S2**).

**Figure 1:**
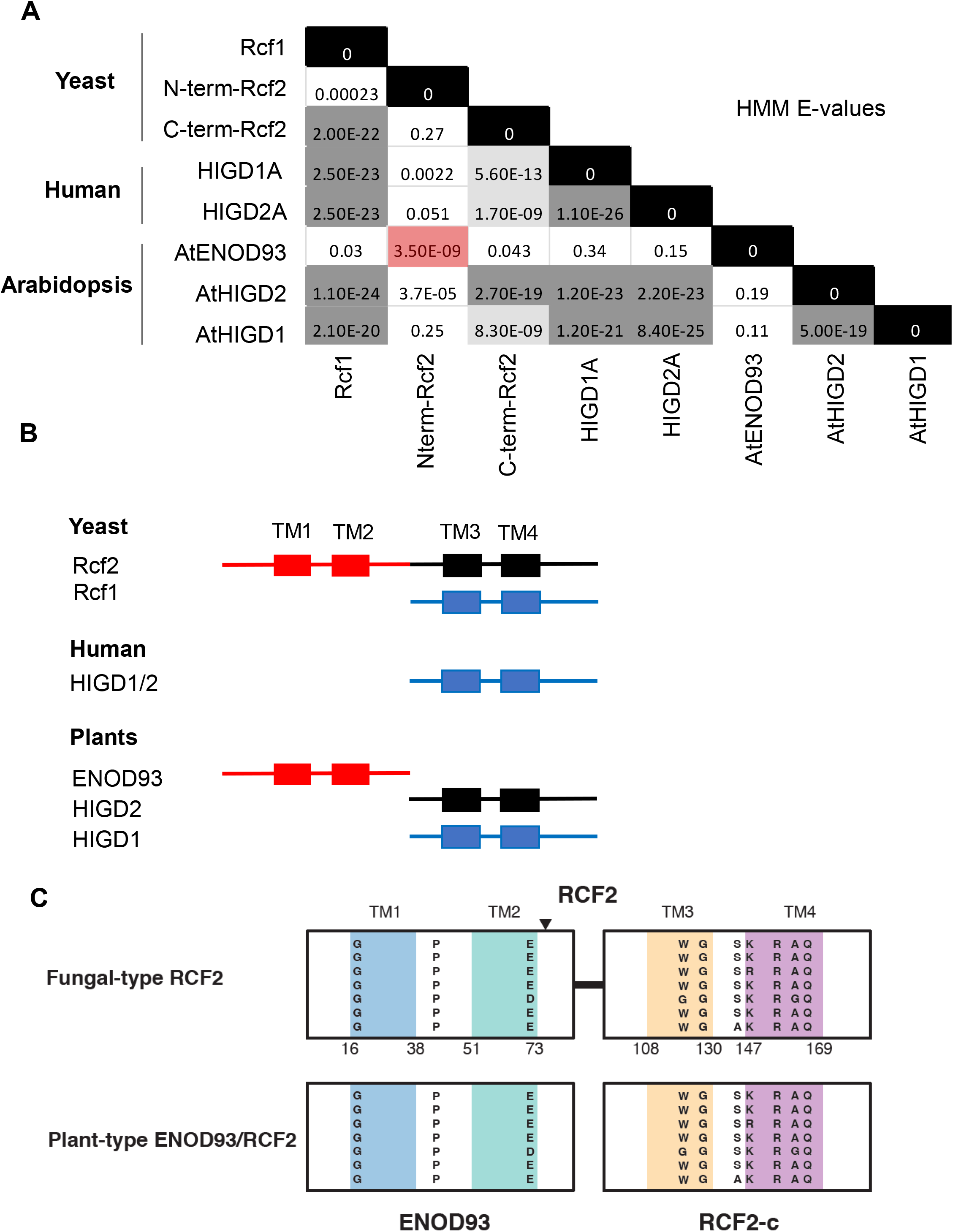
Amino acid sequence similarity between plant, mammal and yeast RCF2 proteins. (A) Hidden Markov model similarities shown as E-values in comparisons of yeast RCF2 N- and C-terminal halves with mammalian HIGD and plant ENOD93 and HIGD proteins. (B) Cartoon of the protein and transmembrane domain structure of RCF, HIGD and ENOD proteins and homology of plant, mammal and yeast proteins with similarity shown by colours.(C) Similarity of transmembrane sequences in TM1-4 between fungal RCF2 and plant-type ENOD93 and RCF2-c like proteins showing single amino acid codes for highly conserved residues at the borders of, and within transmembrane helices.

### Arabidopsis as a genetic model for ENOD93 functional characterisation

To determine if there is a functional similarity between plant ENOD93 and yeast RCF2, given the poor sequence conservation, we used the model plant Arabidopsis to perform genetic and biochemical studies. Arabidopsis encodes a single *ENOD93* gene which co-expresses with many nuclear genes for mitochondrial localised proteins. Of the top 20 most similarly expressed genes (Obayashi et al., 2022), nine are mitochondrial predicted or experimentally mitochondrial located, and five of these are subunits of Complex IV, including the aforementioned ATHIGD2 (**Figure S3A**). To functionally characterise this gene an Arabidopsis mutant *enod93* was isolated from the SALK insertion collection. It contains a T-DNA insertion in its second intron (**Figure 2A**) that fully disrupts *ENOD93* expression (**Figure 2B**). Complementation with an *ENOD93* cDNA in the *enod93* background restored expression. The *enod93* mutant line has a short root phenotype that is complemented in plant lines expressing the *ENOD93* cDNA at 0.1 to 4 times the level observed in WT **(Figure 2B, C**). Growth phenotypes in *enod93* did not extend to the shoot as normal rosette growth is observed in *enod93* and complemented lines (**Figure S4A, C**). Overexpression of *ENOD93* in WT enabled the generation of plant lines with 60 to 100 times more *ENOD93* transcript than WT, but no obvious effect on root or shoot phenotypes (**Figure S4D,E**).

**Figure 2:**
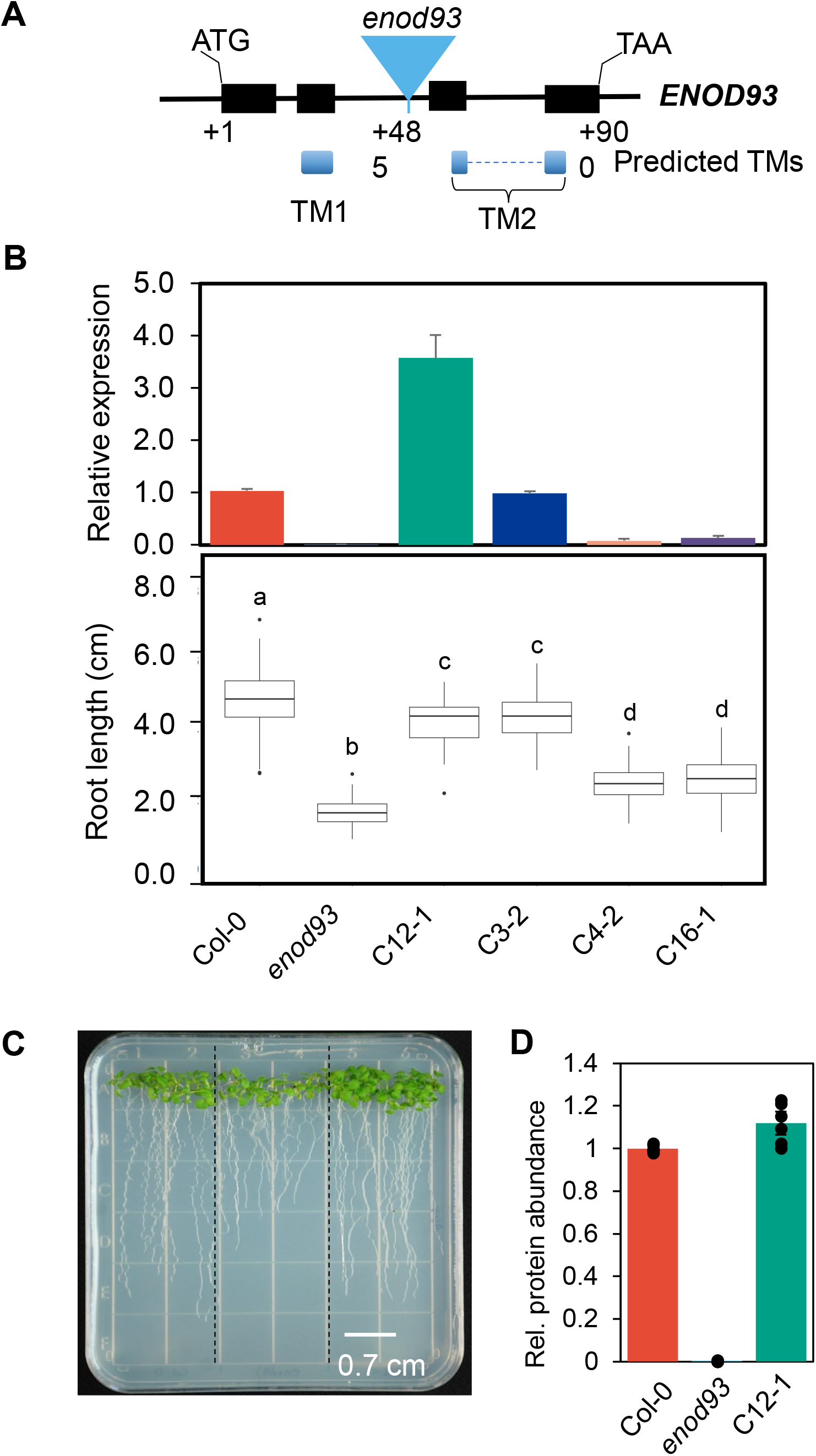
Characterisation of *enod93* mutant and whole plant phenotype in Arabidopsis. (A) Genomic structure of *enod93* t-DNA knockout line showing insertion in intronic region and predicted transmembrane domains based on ARAMEMNON consensus prediction (Schwacke et al., 2003). (B) Expression levels of ENOD93 in different genotypes as determined by qPCR (n = 3). Double asterisks indicate a significant change as determined by Student’s test (**p < 0.01, n = 3). Root length of seedlings of those genotypes is shown below with different letters indicating significant differences based on Kruskal-Wallis rank sum test with a post-hoc Dunn test (P < 0.01, n > 50). (C) Representative image of vertically grown seven-day-old seedlings of Col-0, *enod93* and complemented lines on an agar plate under long day condition (left). (D) Relative ENOD93 protein abundance in WT, *enod93* and C12-1 lines based on targeted LCMS of an *enod93* specific peptide (AAAVAAVAS AIPTVAAVR).

### Localisation of ENOD93 within Arabidopsis mitochondria

To assess the function of ENOD93 we purified mitochondria from plants of WT, *enod93* and a complemented line. To confirm that ENOD93 was absent from mitochondria in *enod93,* we developed a targeted mass spectrometry based MRM assay for an ENOD93 peptide in whole mitochondrial extracts. We showed complete loss of the peptide signal in mitochondrial extracts from *enod93* and recovery to WT levels in the complemented line (**Figure 2D)**. The abundance of a wide series of other respiratory complex subunits and TCA cycle enzymes were also quantified in the genotypes using previously developed MRM assays for specific peptides, but this showed no significant changes in their abundance between WT, *enod93* and complemented lines (**Table S2**).

We also performed native separation of protein complexes by 2D BN-SDS-PAGE (**Figure 2B**). Previously, Klodmann et al (Klodmann et al., 2011) had observed ENOD93 in similar gels as a 12 kDa protein in a large native complex of 250-300 kDa. Native complexome studies from the same group also identified ENOD93 at 250 kDa, and found a small proportion in larger respiratory complexes above 600 kDa and some in the molecular mass region below 100 kDa (Senkler et al., 2017)(**Figure S3B**). We cut out equivalent protein bands from the 250-300 kDa and ∼600 kDa regions of our BN-SDS-PAGE gels and used peptide mass spectrometry to confirm these reported localisations (**Figure S4B**). We identified peptides of ENOD93 in high molecular mass protein complexes above 650 kDa and in the 250 kDa region alongside complex IV subunits as shown previously. In mitochondria isolated from *enod93*, no ENOD93 peptides were identified in the same positions, while we could again detect the presence of peptides for ENOD93 in a complemented line (**Table S1**). This confirmed a co-localisation of ENOD93 with respiratory complexes on native gels in WT and complemented line, and no obvious change in the profile of other BN-SDS-PAGE complexes between WT, *enod93* and the complemented line **(Figure S4B).**

### Loss of ENOD93 leads to progressive loss of respiratory rate in energised Arabidopsis mitochondria

Substrate-dependent oxygen-electrode assays and the addition of cofactors and adenylate nucleotides are typically used to explore functional defects in the mitochondrial respiratory apparatus. We observed a progressive loss of ADP-dependent stimulation of oxygen consumption rate over the course of respiratory assays in mitochondria isolated from *enod93*. During the first addition of ADP and so-called state 3/state 4 transition, *enod93* mitochondria responded similarly to WT. However, following subsequent additions of ADP, stimulation of respiration to the phosphorylating State 3 was not observed (**Figure 3A,B**). This effect was largely reversed in mitochondria from an ENOD93 complemented line. This respiratory effect was independent of the respiratory substrate used in the assay, the same phenomenon was observed in mitochondrial respiration supported by NADH (**Figure 3 A,B**), succinate (**Figure S5 A,B**) or malate and glutamate (**Figure S5 C,D**). To determine if the respiratory rate effect was reversible, we took *enod93* mitochondria that could no longer be stimulated after multiple ADP additions, recovered them from the assay media by centrifugation, and showed they could regain ADP stimulation and that this could be lost again over time by further ADP incubation (**Figure S6)**. We reasoned this process would abolish the membrane potential of the mitochondria and it suggested respiratory capacity could be recovered in this manner. To determine if direct loss of membrane potential could indeed reactivate respiratory rate in *enod93* mitochondria, we added the chemical uncoupler FCCP to respiring *enod93* mitochondria after time-dependent ADP inhibition and found recovery of respiratory rate to that observed in WT and the complemented line (**Figure 3C and D**).

**Figure 3.**
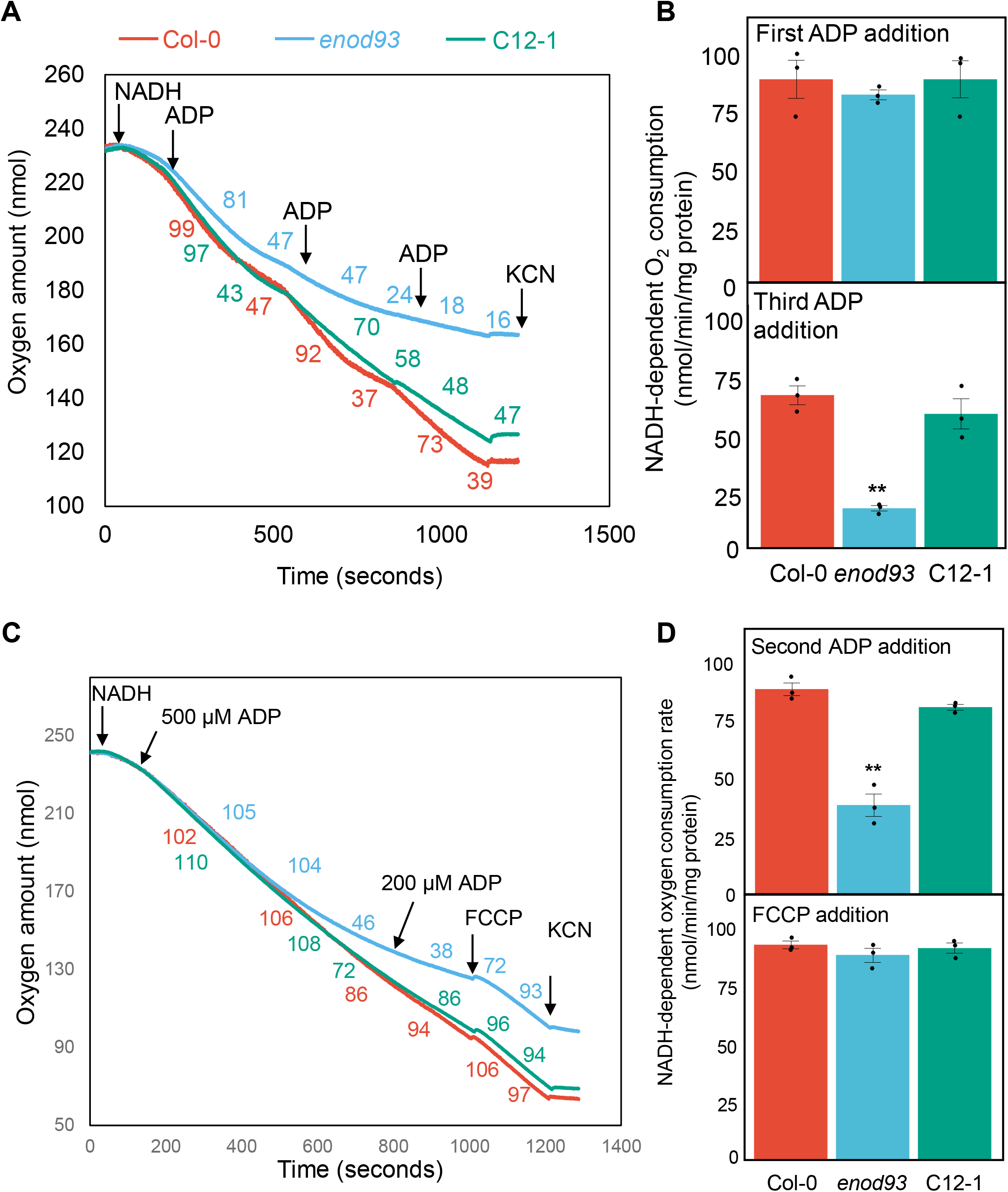
Oxygen consumption characteristics of mitochondria. (A) Representative trace illustrating NADH-stimulated oxygen consumption by mitochondria purified from Col-0, *enod93* and C12-1 seedlings. Oxygen consumption rates in response to the successive treatments are shown in coloured numerical values. Substrate additions to all samples are indicated by arrows with the following concentrations: 10 mM NADH, 0.1 mM ADP and 1 mM KCN. (B) State III oxygen consumption rates for purified mitochondria energized with NADH in response to the first (upper panel) and third (lower panel) ADP addition as indicated in (A). (C) Representative trace illustrating the “recovery” of oxygen consumption rate by isolated *enod93* mitochondria upon FCCP addition. Substrate additions are indicated by arrows with the following concentrations: 10 mM NADH, ADP: 0.2 or 0.5 mM ADP, 10 µM FCCP and 1 mM KCN. (D) State III oxygen consumption rates for purified mitochondria energized with NADH in response to the second ADP addition (upper panel) and to the FCCP addition (lower panel) as indicated in (C). Each data point represents mean ± S.E. with overlaid individual data points as dots (n = 3). Asterisks indicate a significant change as determined by one- way ANOVA with Tukey post-hoc test (* p < 0.05; ** p < 0.01).

### Loss of ENOD93 does not affect respiratory complex abundance or solubilised complex IV activity

To investigate a basis for these effects within the respiratory chain, we followed up the 2D-BN-PAGE analysis **(Figure S4B)** by visualizing the abundance of the major ETC protein complexes in one-dimensional BN-native gels but found no differences in their abundance between genotypes (**Figure 4A**). BN-native in-gel activity assays of complex IV (**Figure 4B**) showed no differences between genotypes, nor did a spectrophotometric assay of complex IV activity in hypotonically-ruptured mitochondrial samples (**Figure 4B**). As there have been reports of ATP inhibition of complex IV in mammalian mitochondria (Acin-Perez et al., 2011; Arnold and Kadenbach, 1997), we directly assessed the effect of ATP on plant complex IV activity (**Figure 4C**). This showed a progressive inhibition of complex IV by increasing ATP concentration in WT, but no differential effect in *enod93* or the complemented line. A small but significant decrease in complex IV activity could only be observed when TMPD + ascorbate was used to directly deliver electrons to cytochrome c which passes them to complex IV in intact mitochondria (**Figure 4D**).

**Figure 4.**
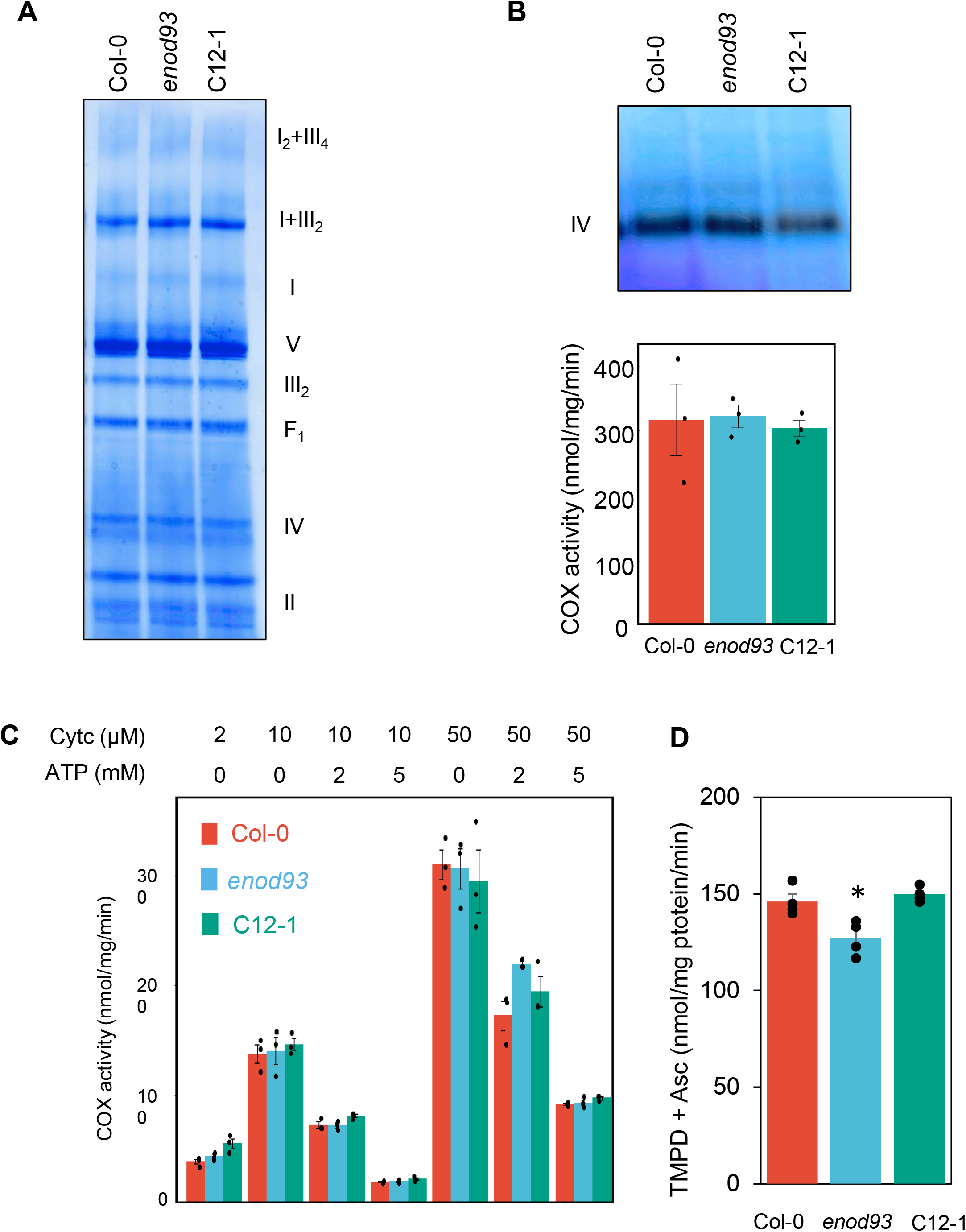
Characterization of mitochondrial respiratory complexes and Complex IV activity in isolated plant mitochondria from *enod93*. (A) Separation of mitochondrial respiratory supercomplexes by 1D-blue-native PAGE. Roman numerals correspond to the locations of respiratory complexes. Gels were visualised by Coomassie Blue. (B) visualization of Complex IV by activity staining of BN-PAGE and cytochrome c oxidase activity in hypotonically-ruptured, freshly isolated mitochondria in the presence of 50 µM cytochrome c. No significant change was found based on one-way ANOVA (n = 3). (C) Sensitivity of cytochrome c oxidase activity by in hypotonically- ruptured, freshly purified mitochondria in the presence of different cytochrome c and ATP concentrations. No significant change was found between genotypes for each treatment based on one-way ANOVA (n = 3). (D) ADP dependent respiratory rate with TMPD +Asc as substrates in intact plant mitochondria. All data represents mean ± S.E. with overlaid individual data points as dots (n = 3).

### Loss of ENOD93 slows ATP synthesis and raises mitochondrial membrane potential

Loss of ADP-stimulated State 3 respiration rate (**Figure 3, Figure S4**) suggested a progressive loss of ATP synthesis rate was occurring in *enod93*. To prove this, we directly assayed ATP synthesis and ADP depletion rate in energized mitochondria. We show there was a marked impact in *enod93* mitochondria on the rate of ADP use and ATP generation over time (**Figure 5A**). To determine if the reverse rate of ATP hydrolysis by the mitochondrial F_1_F_o_ ATP synthase was also affected, we measured it spectrophotometrically in broken mitochondria and showed it was unaffected in *enod93* (**Figure 5B)**. Analysis of membrane potential during oxidative phosphorylation using safranin fluorescence showed that *enod93* mitochondria maintained a higher membrane potential throughout respiratory assays than WT or complemented lines, suggesting a restriction of respiratory rate was occurring. In all genotypes the addition of FCCP depleted the membrane potential (**Figure 5C)**, and by inference from respiration rate studies **(Figure 3C,D)** can then reactivate complex IV activity. To confirm the importance of this elevated membrane potential on the effect in endo93 mitochondria, we assessed the effect of malonate addition during succinate dependent respiration and showed when membrane potential was decreased there was little difference between WT and enod93 in ADP dependent oxygen consumption rate (**Figure S7A,B**). To show that the operation of complex IV was essential for the elevated membrane potential in enod93 mitochondria we repeated the experiment in **Figure 5C** using pyruvate and malate as substrate to allow proton translocation via Complex I and III but provided ferricyanide as an artificial electron acceptor to bypass complex IV. When complex IV was bypassed in this way there was no longer any difference in membrane potential between WT, *enod93* and complemented line mitochondria (**Figure S7C,D**).

**Figure 5:**
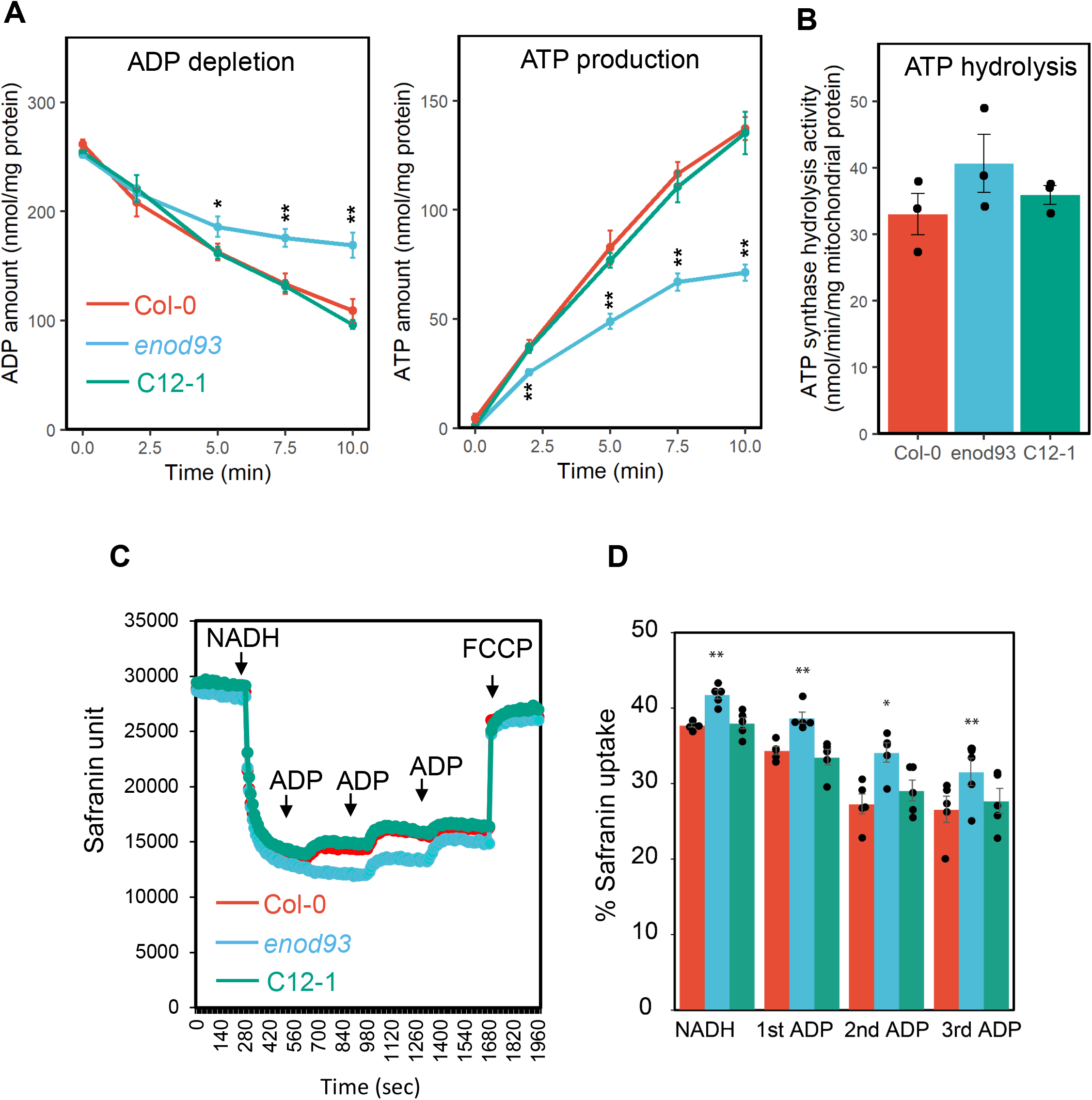
Forward and reverse modes of ATP synthase activity and membrane potential in isolated *enod93* mitochondria. (A) Time course of ADP depletion (left) and ATP production during NADH- dependent respiration by purified mitochondria. Reaction was initiated after the addition of NADH (10 mM) and ADP (1 mM) to 20 μg of freshly isolated mitochondria and terminated by excess amount of ice-cold 15% TCA solution. Following metabolite extraction and solid phase extraction, ADP and ATP concentrations were determined by LC-MS. Each data point represents mean ± S.E. (n = 3), and asterisks indicate a significant change between Col-0 and *enod93* and between *enod93* and C12-1 as determined by one-way ANOVA with Tukey post-hoc test (* p < 0.05; ** p < 0.01). (B) ATP hydrolysis rate of broken mitochondria. Mitochondrial fraction (20 μg) was subjected to repeated freeze-thaw cycles before 1.5 mM ATP was added to initiate the hydrolysis reaction. ATP hydrolysis rates were determined by monitoring the production of ADP over time using LC-MS. Data represents mean ± S.E. with overlaid individual data points as dots (n = 3). No significant change was found based on one-way ANOVA. (C) Representative trace of safranin fluorescence as a measure of membrane potential in mitochondria from WT, *enod93* and complemented line after multiple ADP additions during NADH-dependent respiration. (D) Difference in percentage safranin uptake after 1^st^, 2^nd^ and 3^rd^ ADP addition in isolated endo93 mitochondria (mean ± S.E. with overlaid individual data points as dots n = 4).

### Loss or overexpression of ENOD93 alters oxidative phosphorylation and metabolism in whole plants

To determine if the effect in *enod93* isolated mitochondria could be observed in intact plants we conducted a series of extra experiments based on our findings. We found whole seedling respiration rate was elevated in *enod93* seedlings but not in an ENOD93 complemented line (**Figure 6A**), but there was no difference in respiratory rate responses to the addition of the membrane uncoupler FCCP amongst genotypes (**Figure 6B**). Additionally, we found that the root growth phenotype of *enod93* was more sensitive to FCCP addition to the growth medium than WT or complemented lines, suggesting a longer-term sensitivity in *enod93* seedlings to reduced ATP production rate in intact roots (**Figure 6C**).

**Figure 6:**
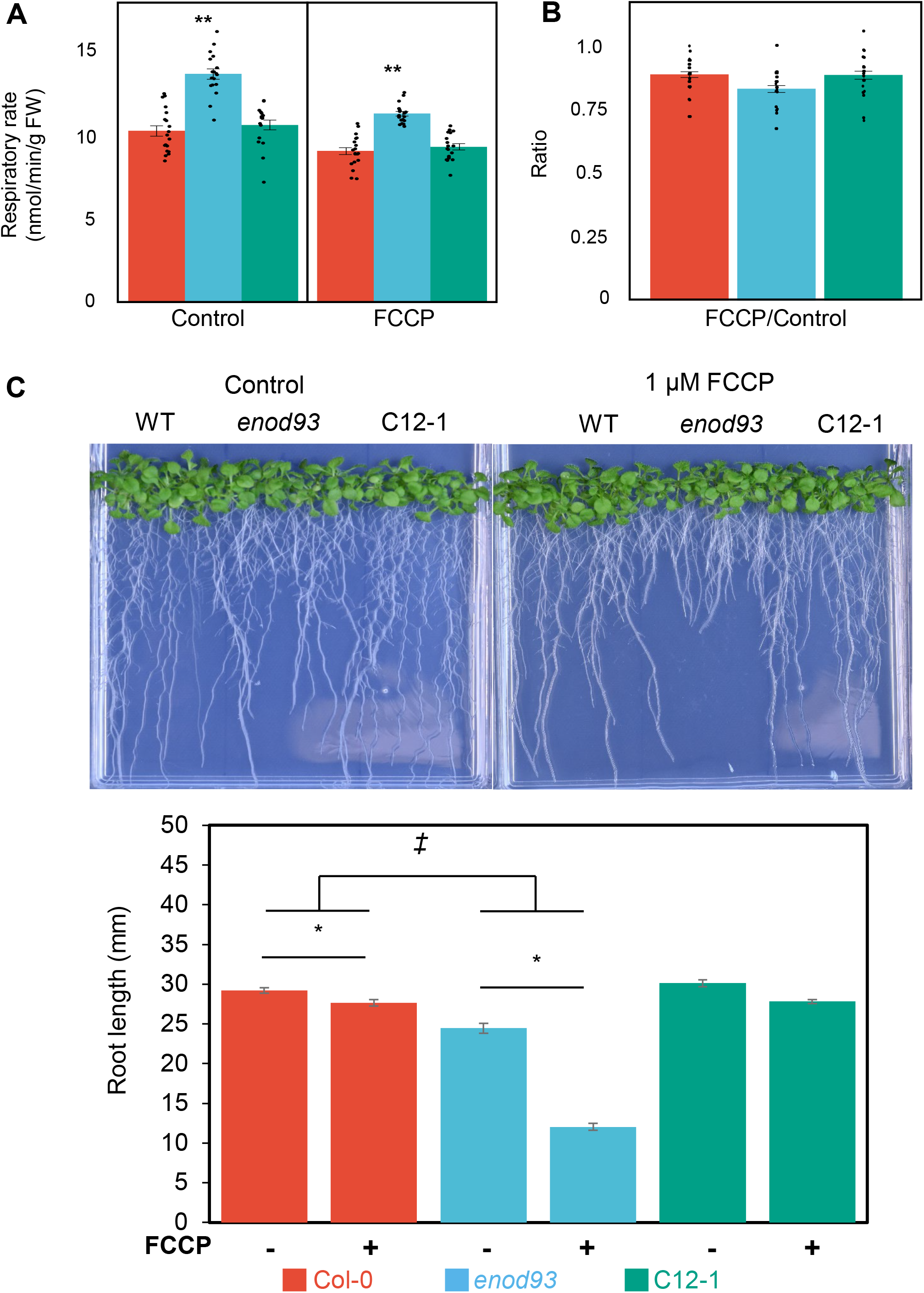
Effect of *enod93* loss and FCCP addition on whole plant respiration and root growth. (A) Basal respiration rates of seven-day-old Arabidopsis seedlings were measured. FCCP (2 μM) was then added and allowed to equilibrate for 20 min before rates were measured (right). Asterisks indicate a significant change as determined by one-way ANOVA with Tukey post-hoc test (** p < 0.01, n ≥ 9). (B) Ratio of basal respiration rates to FCCP-stimulated respiration of seedlings (n ≥ 9). No significant change was found based on one-way ANOVA. All measurements were carried out at 25°C using a Clarke-type oxygen electrode. Data represents mean ± S.E. with overlaid individual data points as dots. (C) Representative image of the effect of FCCP on root growth in enod93, Col-0 and C12-1 line. Bar graph of the differences in root length between enod93 and Col-0. Asterisks represent significant treatment effects, double daggers represent significant genotypic effects (two-way ANOVA; n ≥ 17; p < 0.01).

To directly assess adenylate levels in whole tissues, root samples from a range of genotypes were snap frozen and extracted to measure nucleotide levels. The whole root ADP content was found to be nearly twice as high in *enod93* than WT without a change in root ATP content. Complementation with ENOD93 to different degrees progressively lowered ADP content to WT levels **(Figure 7A,B)**. The ATP/ADP ratio approached 6 in WT, lowered to below 4 in *enod93*, and recovered in ENOD93 complemented lines, but was found to decrease to even lower levels in ENOD93 overexpression lines (**Figure 7C**).

**Figure 7:**
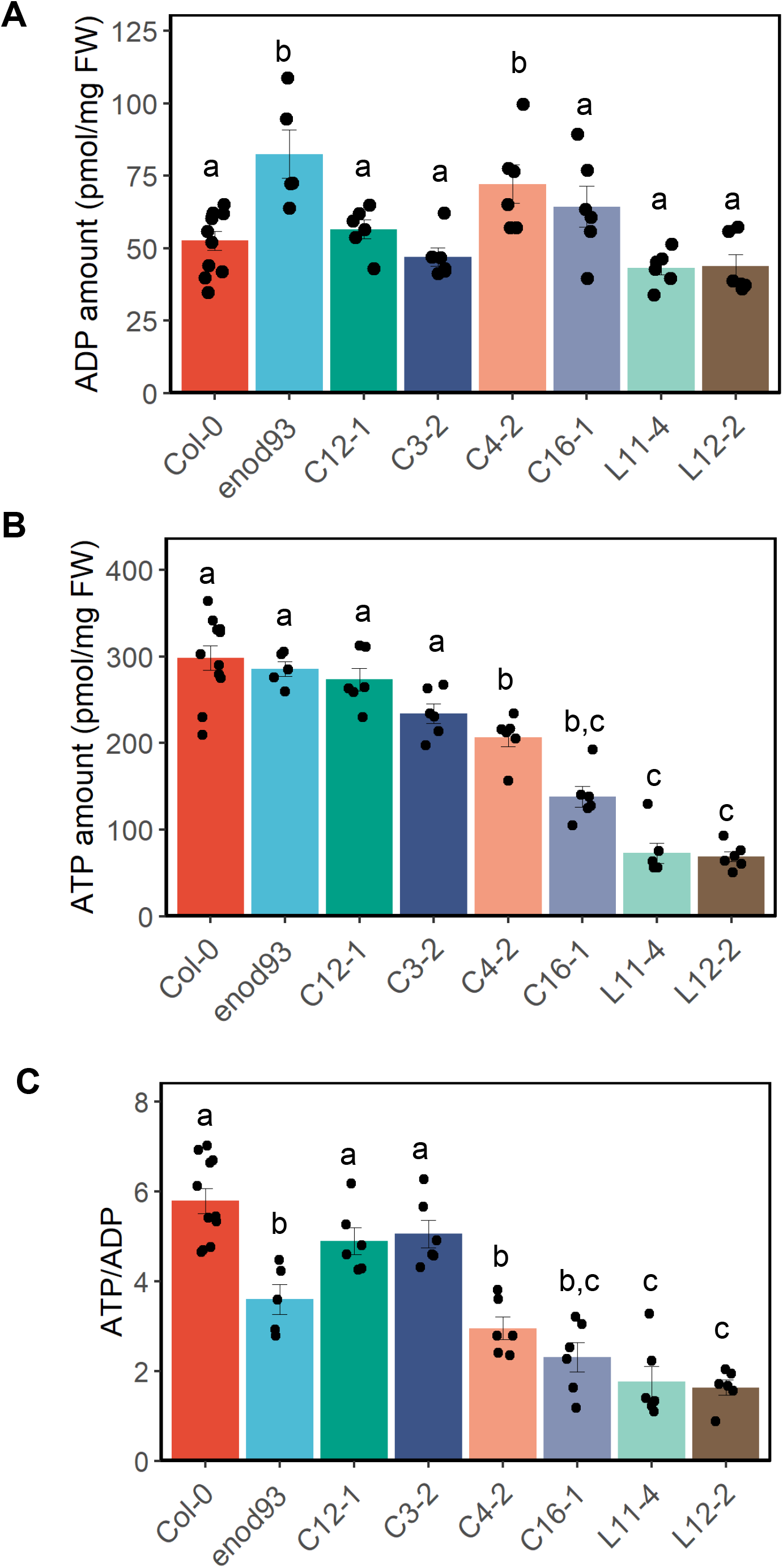
Quantitation of ADP and ATP in roots from WT, enod93, complemented and over- expression genotypes. Absolute concentrations of ADP (A) and ATP (B, in pmol/mg FW) and the corresponding ATP/ADP ratios (C) were quantified in roots from a pool of seven-day-old seedlings. Data shown are mean with overlaid individual data points as dots (n ≥ 5). Different letters indicating significant differences based on a one-way ANOVA test with a Tukey post-hoc test (P < 0.05). *enod93* complemented lines C12-1, C3-2 and C4-2 are shown. Overexpressed *ENOD93* lines L11-4 and L12-2 are shown.

Overexpression of OsENOD93-1 in rice had previously been shown to increase amino acid content (Bi et al., 2009). A selection of the Arabidopsis genotypes with altered energetic states (**Figure 7**) was therefore profiled for their absolute abundance of a range of amino acids and organic acids in shoots and roots. These data showed elevation of alanine, glycine, fumarate and succinate levels in *enod93* roots, compared to WT levels, and recovery of the same organic and amino acid levels in complemented lines (**Figure 8, Figure S8**). In contrast, ENOD93 overexpression lines had increased leucine and isoleucine levels above WT levels in roots (**Figure 8**), and increased histidine, proline and valine above WT levels in leaves (**Figure S9**).

**Figure 8:**
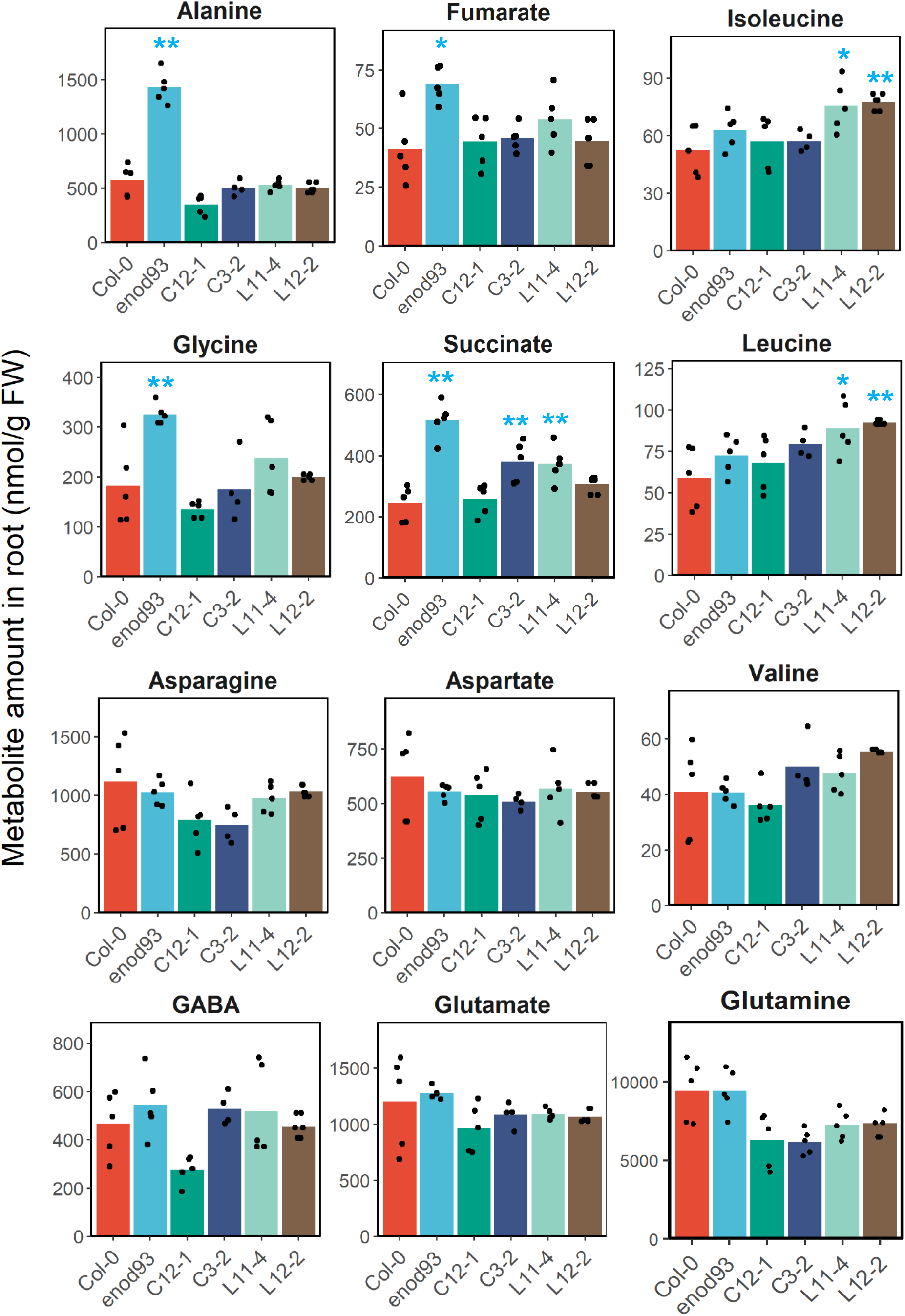
Analysis of amino acids and organic acids in roots from different ENOD93 genotypes. Roots excised from a pool of seven-day-old seedlings were collected in liquid nitrogen and extracted metabolites analysed by LC-MS. Metabolites that demonstrated significant differences between genotype(s) or involved in nitrogen metabolism are shown, and the remaining metabolites are shown in Supplementary Figure S8 and S9. Data from *enod93* complemented lines C12-1 and C3-2 and overexpressed *ENOD93* lines L11-4 and L12-2 are shown. Data shown are mean with overlaid individual data points as dots (n ≥ 5) in nmol per gram fresh weight (FW), and asterisks indicate a significant change relative to Col-0 as determined by Student’s t-test (* p < 0.05; ** p < 0.01).

## Discussion

ENOD93 is a small, ubiquitously expressed plant membrane protein with an unknown protein domain that has evaded functional evaluation for decades (Bi et al., 2009; Kouchi and Hata, 1993; Reddy et al., 1998; Yan et al., 2015). In its absence, we show that plant mitochondria suffer a progressive loss of function which to our knowledge has never been reported for any other mutant of an ETC component in plant mitochondria. We also show that adenylate levels and metabolic pathways are perturbed *in vivo* in the absence of ENDO93 leading to reduced root growth. Combining evidence from genetics, mitochondrial biochemistry and phylogenetic sequence analysis we show ENOD93 likely plays a widespread and ancient role in ensuring the function of the mitochondrial electron transport chain by interacting with Complex IV and influencing its function in energized mitochondria.

### ENOD93 affects plant mitochondria function and is associated with Complex IV

Loss of ENOD93 has a profound impact on respiratory function and ATP synthesis in Arabidopsis, however, precisely explaining the mechanism of this phenomenon remains complex. We show this effect is independent of the respiratory substrate used in the oxygen consumption assays, indicating the lesion is downstream of respiratory substrate dehydrogenases. We observe the effect of ENOD93 loss on total CIV activity is minor, only becoming evident when it is operating as part of the respiratory chain in energized mitochondria. The loss of ENOD93 slows the oxygen consumption rate to a much greater degree when the membrane potential has increased to a certain level and this is then linked to a marked decrease in ATP synthesis rate. The importance of a high membrane potential in the reversible inactivation of Complex IV and ATP synthesis in *enod93* in underlined by the two scenarios that appear to lead to recovery. Firstly, we show centrifugal recovery of mitochondria and re-assaying of oxygen consumption that removes respiratory substrates and de- energize the membrane allows recovery (**Figure S5**) and secondly we evidence that addition of FCCP that will dissipate the inner membrane electrical potential recovers full Complex IV activity **(Figure 3)**.

The evidence for a physical link between ENOD93 and Complex IV comes from a series of published proteomics data, the case of the related RCF2 protein from yeast, and the information provided in this report. Gel-spot identifications from BN-PAGE separations of plant mitochondria first observed a native complex of ∼250 kDa that contained peptides for ENOD93 (Klodmann et al., 2011) and also showed that HIG2 is associated with CIV (Millar et al., 2004). Subsequently, complexome profiling of similar gels tracked quantitative changes in ENOD93 abundance and again found it in a mass range equivalent to other CIV subunits (Senkler et al., 2017), but this association was not claimed at the time due to the significant proportion of ENOD93 being found in a dissociated state towards the bottom of BN-PAGE separations. Re-clustering of these data now shows that both ENOD93 and HIG2 are in the protein complex with CIV subunits and that the ENOD93 in lower native mass complexes is associated with Cox6b **(Figure S2B)**. In yeast, RCF2 was not found in early crystal structures of CIV, but more recent CryoEM CIII/CIV super complex structures (Hartley et al., 2020; Moe et al., 2023) contain the C-terminal HIG domain of RCF2 bound to Cox12 (which is the yeast equivalent of COX6b in plants). CryoEM of the plant complex IV from mung bean did not identify HIG2 or ENOD93 in the structure but did observe HIG2 by mass spectrometry in samples used for the structural determination (Maldonado et al., 2021); consistent with the instability of this association. The RCF2 association with the CIV2/CIII supercomplex in CryoEM structures is far away from the CIV/CIII interface, suggesting it is definitively with each CIV molecule separately. Collectively this information is consistent with HIG2 and ENOD93 both being associated with intact Complex IV in plants and ENOD93 dissociating from the complex in detergent preparations in association with Cox6b, providing a putative site of Complex IV interaction.

### ENOD93 is similar to the N-terminus of RCF2 and they share a degree of functional equivalence

Evaluating the evidence for functional equivalence of RCF2 and ENOD93 is complicated because ENOD93 is only homologous to the N-terminal domain of RCF2. Much of the evidence for the functional role of RCF2 in yeast arises from a study of Δrcf1, Δrcf2 or Δrcf1rcf2 mutants. Loss of RCF1 decreases the abundance of complex IV and it appears to have a role in CIV assembly through binding to COX3 (Strogolova et al., 2012). In contrast, loss of RCF2 does not change complex IV abundance, but alters the activity of the enzyme (Strogolova et al., 2019). Complementation studies in yeast show that RCF1 and RCF2 are not functionally redundant (Homberg et al., 2021; Rompler et al., 2016). Yeast mitochondrial lacking both RCF proteins have lower mitochondrial membrane potential and the respiratory chain becomes inactive over time (Strogolova et al., 2019). In mutants lacking both RCF1 and RCF2 proteins there is a proton leak back through complex IV, suggesting that the HIG domains of RCFs might be involved in preventing proton leak from the IMS to the matrix through complex IV (Hoang et al., 2019). Recently it was also found that fusing the RCF2 N-terminus to RCF1 could rescue an Δ*rcf2Δrcf3* mutant (Homberg et al., 2021) which was surprising as previously the focus had been only on the RCF2 C-terminal HIG domain as having unique capabilities in the regulation of complex IV activity (Rompler et al., 2016).

Based on our phylogenetic and functional evidence we propose that RCF2 in yeast is a fusion of two genes found in other eukaryotes that may have related but different functions, which complicates the interpretation of joint loss experiments and the chimeric forms of RCF2 that have been generated and studied. Independent function of the RCF2-N terminal and RCF2-C terminal is certainly possible given that yeast RCF2 can be proteolytically cleaved inside mitochondria into these two parts, allowing its two halves to function independently at a biochemical level (Rompler et al., 2016). Collectively, (a) the functional evidence that yeast RCF2 functions in CIV activity regulation associated with the protonmotive force and the related observations in *enod93* plants, (b) the physical association of RCF2 C-terminus with COX12 in the CryoEM structure and the co-migration of ENOD93 and COX6b in plants, and (c) the profile hidden Markov model link between the RCF2 N-terminus and ENOD93 in phylogenetic studies across eukaryote, provide a substantial basis for suggesting a functional equivalence between these two proteins. In addition at the whole cell level, Δrcf2 elevated whole yeast cell respiration rate (Strogolova et al., 2019) as observed in *enod93* plants (**Figure 6A**), and yeast Δrcf2 cells were more sensitive to the K+/H+ ionophore nigericin (Strogolova et al., 2019) as *enod93* plants were to the uncoupler FCCP (**Figure 6B**). One notable difference is that the progressive respiratory inhibition in *enod93* in isolated plant mitochondria appears to be fully reversible by membrane de-energization, while the progressive respiratory inhibition observed in yeast Δrcf1Δrcf2 is considered to be an irreversible complex IV suicide inactivation requiring resynthesis of the enzyme (Hoang et al., 2019). This difference may underlie distinct roles of the N and C terminal domains of RCF2 that have yet to be functionally evaluated in yeast by deleting only the N-terminus of RCF2 and studying mitochondrial functions.

### ENOD93 function is linked to nitrogen fixation and nitrogen metabolism in plants through ATP demand

ENOD93s were previously identified in plants as genes with elevated expression in N_2_ fixing root nodules and/or highly responsive to plant nitrogen status (Bi et al., 2009; Cabeza et al., 2014; Kouchi and Hata, 1993; Yan et al., 2015). In N_2_ fixing root nodules, mitochondria in infected cells operate at a low oxygen concentration and an elevated membrane potential to generate high rates of ATP synthesis to sustain the symbiosis (Millar et al., 1995). Based on our discovery of ENOD93 function, the nodule enhanced expression of ENOD93 and the ability of miR393j-3p to regulate it and significantly influence nodule formation in soybean (Yan et al., 2015) can be reinterpreted as a biochemical effect of this gene on ATP availability. Cabeza et al also showed there is a strong correlation between nitrate inhibition of nodule N_2_ fixation in Medicago and downregulation of *ENOD93* (Cabeza et al., 2014). Close inspection of their supplemental transcript profiling data also shows this effect is correlated with downregulation of other complex IV subunit genes, suggesting mitochondrial inhibition of complex IV is responsible. The biotechnological use of ENOD93 to alter nitrogen use efficiency in rice (Bi et al., 2009) is consistent with the fact that nitrate uptake and assimilation both use a substantial proportion of cytosolic ATP in plants (Glass, 2003; Xu et al., 2012). The specific cost for net NO_3_^-^ uptake ranges from 3 to 5 ATP in plant species growing at different rates (Scheurwater et al., 1999). Following reduction to ammonia, assimilation of N into amino acids also has a substantial additional ATP cost via glutamine synthase (GS) and asparagine synthase (ASN). In *enod93*, the two amino acids that accumulate the most are Ala and Gly (**Figure 8**) which are the lowest cost AAs to make based on metabolic cost calculations (average cost 23 ATP/AA; (Arnold et al., 2015)). In contrast the AAs that accumulate in ENOD93 OE lines (Ile and Leu in roots, and His, Pro and Val in leaves) are products of pyruvate, oxaloacetate and 2-oxoglutarate, and cost 57 ATP/AA on average to build (Arnold et al., 2015). Therefore, the changes in metabolite levels recorded are consistent with a limitation in respiration rate in *enod93* and an accumulation of low ATP costs AAs, while following ENOD93 overexpression an accumulation of higher cost AAs is recorded. The apparent N control of ENOD93 expression makes it a sensible system to enable plants to modulate ATP synthesis rate to sustain N uptake and assimilation of amino acids and provide the high ATP demand needed in N_2_ fixing root nodules. If ENDO93, like RCF2, is a supernumerary subunit of Complex IV then changes in its abundance by altered expression patterns will render a different proportion of Complex IV molecules under its control allowing a tuning of ATP synthesis.

### ENOD93 is an ancient and widely distributed component of mitochondria

The phylogenetic distribution of ENOD93 across green plants and in iterative searches of the NCBI nr database using psi-blast showed this domain is very widely observed in many eukaryotic lineages (**Figure S10**). ENOD93 homologs, can be consistently identified in all major groups interrogated, including green algae, and early-diverging plant lineages (*e.g.*, hornworts, mosses, lycophytes, ferns), along with diverse gymnosperms and angiosperms. Using a profile Hidden Markov model based on green plant ENOD93 to interrogate protein sequences from diverse microbial eukaryotes, we were also able to identify putative highly divergent ENOD93 homologs in multiple non-plant eukaryotic lineages, including cryptophytes, amoebozoans, collodictyonids, apusomonads, diverse stramenopiles, haptophytes, and basal holozoans (*i.e.*, fungi and unicellular relatives of fungi and animals, but not animals themselves), though homologs were not identified in many other groups **(Figure S10)**. Although functional annotation is not available for any of these putative ENOD93 homologs, candidate homologs from amoebozoans *Acanthamoeba castellanii* and *Dictyostelium discoideum* have been identified in highly purified mitochondria (Freitas et al., 2022; Gawryluk et al., 2014). The patchy but very wide distribution of putative ENOD93 homologs across diverse lineages suggests its ancient origins within eukaryotes. The uniting of functional evidence for its role in plants and yeast shown here provides a strong foundation for its wider evaluation across eukaryotes as an important factor in mitochondrial ATP synthesis regulation.

## Materials and methods

### Homology detection

To investigate the distribution of ENOD93 homologs across land plants and green algae, the *Arabidopsis* homolog (AT5G25940.1) was used as a query in iterative psi-blast searches of the NCBI nr database (Altschul et al., 1997). After each iteration, evident ENOD93 homologs from diverse lineages, including charophytes, zygnematophytes, liverworts, hornworts, mosses, ferns, lycophytes, and angiosperms, were manually selected to refine the position-specific scoring matrix in subsequent searches. Putative ENOD93 homologs were identified in other eukaryotic groups using profile Hidden Markov model (HMM) searches of phylogenetically broad protein sequence datasets including MMETSP (Keeling et al., 2014) and PhyloFisher (Tice et al., 2021), along with select individual datasets. Initial searches were performed using Hmmer v3.3.2 based on multiple alignments of diverse plant and green algal ENOD93 homologs generated using MAFFT v7.475 under the L-INS-i iterative refinement method (Rozewicki et al., 2019). Where likely homologs of ENOD93 were identified in other eukaryotes, those homologs were added to the initial alignment, from which a broader HMM was generated and again used to iteratively query the datasets described above. Significant matches were defined as those that met the default Hmmer inclusion threshold and reuse. In order to increase our confidence in assigning plant ENOD93 and Rcf2 proteins as *bona fide* homologs of the fungal RCF2 N- and C-terminal regions, respectively, we also tested the reciprocal HMM search, with the expectation that fungal RCF2 queries retrieve homologs of both plant ENOD93 and RCF2. Briefly, homologs of RCF2 from diverse fungi were collected and aligned as above and used to generate profile HMMs and to query plant datasets. Phylogenetically broad bioinformatic searches for homologs of RCF1 and RCF3 were carried out essentially as above, though the high conservation of RCF1 precluded the need for HMM searches. Prediction of transmembrane helices was done with DeepTMHMM v1.0.19 (Hallgren et al., 2022). However, even known transmembrane helices (*e.g.*, in yeast RCF2) were not predicted accurately. In cases where the expected number of transmembrane helices was not predicted, Kyte-Doolittle hydropathy plots were manually inspected for putative ENOD93 and RCF2 sequences to verify their physicochemical plausibility as homologs. The E-values for HMM pair-wise comparisons shown in Figure 1B were generated using HHpred (Zimmermann et al., 2018).

### Plant material and growing conditions

A seed stock of an Arabidopsis T-DNA insertion mutant line (SALK_204202; in Columbia-0 background, and herein *atenod93*) was identified and obtained from the Arabidopsis Biological Resource Center (ABRC). When planted on soil, seeds were stratified for 48 hours and then spread on soil. Plants were grown in a growth chamber maintained under a 16-hour photoperiod (100-150 μmol m^-2^ s^-1^) at 21°C during the day and 18°C during the dark, with 60% relative humidity. For all *in vitro* experiments, plants were grown on agar plates using 1% agar supplemented with ½-strength Murashige & Skoog (MS) basal salts (PhytoTech Labs) at pH 5.7, 1% sucrose, and without additional nitrogen sources. Seeds were sterilized by shaking them for 10 mins in 70% ethanol and 0.05% Triton-X. The solution was removed and seeds were allowed to sit for 5 mins in 100% ethanol and then placed in fresh ethanol for an additional 3 min. Seeds were allowed to dry on sterile filter paper in a flow hood and kept in sterile microfuge tubes. After placing them on plates, they were allowed to stratify at 4°C in the dark for 48 hours. Unless stated otherwise, plates were grown vertically under a 16-hour photoperiod as described above. To isolate mitochondria, surface-sterilized seeds were grown in ½-strength MS medium, 2 mM MES pH 5.7) supplemented with 1% (w/v) sucrose and 0.1% (w/v) agar for 14-16 days with gentle agitation (40-60 rpm) under long day conditions. For FCCP sensitivity of root phenotypes, seeds are surface sterilized and stratified as described above. Arabidopsis seedlings were grown for two weeks on vertical one-half strength Murashige and Skoog agar (10 g·L^-1^ agar, 10 g·L^-1^ sucrose, 0.4 g·L^−1^ MES) plates with and without 1 µM FCCP added before autoclaving. 20 mM FCCP stock was made in 100% ethanol. Root lengths were measured and analysed using ImageJ 1.52v.

### Generation of AtENOD93 overexpression and complementation lines

The nucleotide sequence of the *A. thaliana ENOD93* gene was retrieved from the NCBI database with the GenBank ID At5g25940. The full-length coding region of AtENOD93 was amplified from Arabidopsis cDNA using the primer pairs AtENOD93-GFwd and AtENOD93-GRev to generate a construct for overexpression in wild-type Col-0 (WT) and *atenod93* mutant backgrounds employing the Gateway technology with pDONR™ 221 as the entry vector and pB2GW7 as the destination vector. The resultant 35S:AtENOD93 construct was amplified in *E. coli* DH10b cells and then transferred to *Agrobacterium tumefaciens* strain LBA4404 which was used to transform wild-type (Col-0) and mutant plants via the standard floral dip method (Clough and Bent, 1998) and screened with 0.1% solution of the herbicide Basta^®^ (glufosinate; Bayer CropScience). Positive transgenic lines were grown to T2 and T3 generations to obtain homozygous seeds and seeds from the T3 generation were used in subsequent studies.

### Genotyping of the transgenic and T-DNA insertion lines

The putative transgenic plants that survived the herbicide treatment were confirmed for the integration of the transgenes by PCR using genomic DNA as a template. DNA samples were obtained using the TPS extraction method (Edwards et al., 1991). The PCR reactions were run with Platinum Taq DNA Polymerase (Life Technologies) and using a vector-specific primer (pB2GW7-Fwd) and a promoter-specific (35S-Fwd) primer, either combined with the transgene-specific primer AtE93-Rev (listed in Supplementary Table S3).

For verifying the T-DNA insertion site in *atenod93*, wild-type and *atenod93* lines were genotyped using primers designed via SALK’s iSect Primer program (atenod93-LP, atenod93-RP, and LBb1.3). To sequence the left border of the T-DNA insert and the adjacent genomic sequence, the region was first amplified using the primers TDNA-SeqLB-R and LBb1.3. The resulting amplicon was then sequenced using the same primers used for the amplification. All primers used for cloning and genotyping are listed in Table S3.

### Quantitative RT-PCR analysis

To analyse the expression of *AtENOD93*, total RNA was isolated from 100 mg of rosette leaf tissue from WT, *atenod93*, overexpression, and complementation lines using the Total Plant/Fungal RNA Isolation Kit (Norgen Biotek Corp). After removing any contaminating DNA, 1 µg of the total RNA from each sample was reverse transcribed into cDNA using qScript™ cDNA SuperMix (Quanta Biosciences) according to the manufacturer’s protocol. Real-time RT-PCR was performed in a 20 µl reaction using 50-100ng of cDNA from each sample and the PerfeCTa® SYBR® Green SuperMix (Quanta Biosciences) on the ABI7300 system (Applied Biosystems). The PCR program adjusted for the amplification of the target fragment consisted of one cycle at 95°C for 2 min, and 40 cycles of 95°C for 15 s and 60°C for 30 s. The specific primers AtENOD93-qFwd and AtENOD93-qRev were designed with the software Primer3 and used to amplify *AtENOD93* cDNA. Relative expression values from two technical and three biological replicates were calculated using Applied Biosystem’s included software, which uses the 2^-ΔΔCT^ method. The *Actin7* gene was used as the internal reference for normalization. Primers used in expression analysis are listed in Table S3.

### Isolation of mitochondria, and measurements of oxygen consumption,, ATP synthesis, and membrane potential

Mitochondria were isolated from two-week-old Arabidopsis seedlings as described previously (Sweetlove et al., 2007). Substrate-dependent O_2_ consumption by purified mitochondria was measured in a computer-controlled Clark-type O2 electrode unit according to Lee et al. (Lee et al., 2010), using 1 ml of respiration medium (0.3 m sucrose, 5 mm K_2_H_2_PO_4_, 10 mm TES, 10 mm NaCl, 4 mm MgSO_4_, 0.1% (w/v) BSA, pH 7.2) and 100 μg of mitochondrial sample. Cytochrome c oxidase activity was determined by the rate of oxygen consumption (Neuburger et al., 1982), except different concentrations of exogenous cytochrome c and ATP were added. For measurement of O_2_ consumption by cytochrome c oxidase activity with endogenous cytochrome c, 300 µM N,N,N’,N’- Tetramethyl-p-phenylenediamine dihydrochloride (TMPD) plus 10 mM ascorbate were added.

Substrate-dependent forward mode of ATP synthase activity (i.e. ATP production) was determined by incubating 20 µg freshly isolated mitochondria in 200 µl respiration medium with 0.1 mM NADH and 1 mM ADP. The reaction was terminated at the specified time by adding 800 µl 15% trichloroacetic acid and snap-freezing in liquid nitrogen. For the reverse mode of ATP synthase activity (i.e. ATP hydrolysis), isolated mitochondrial fractions were subjected to at least three freeze-thaw cycles in liquid nitrogen. Following resuspension in respiration buffer, mitochondria (20 µg in 200 µl) were incubated in 1 mM ATP. At the specified time, the reaction was stopped by adding 800 µl 15% trichloroacetic acid and snap-freezing in liquid nitrogen. ADP and ATP concentrations in these samples were determined by mass spectrometry described below.

Safranin O was used as a membrane-permeable cation that crosses to the matrix surface of the inner mitochondrial membrane in proportion to the density of negative charge. Membrane potential across the inner mitochondrial membrane was determined by the decrease in safranin O fluorescence. For each assay, 20 µg freshly isolated mitochondria were incubated in 200 µl respiration medium containing 1 µM safranin O. Mitochondria were equilibrated for 5 minutes before state 2 respiration was initiated by respiratory substrate addition. State 3 respiration was established by multiple additions to the final concentration of 1 mM ADP. The reaction was terminated by adding 4 µM FCCP to dissipate ΔΨ to zero. Safranine O fluorescence was recorded using FLUOstar® Omega plate reader (excitation at 544 nm and emission at 590-10 nm). The percent safranin uptake at any time point was calculated as (micromolar safranin at 0 ΔΨ − micromolar safranin at time x)/micromolar safranin at 0 ΔΨ.

### Blue native electrophoresis and protein identification by liquid chromatography tandem mass spectrometry (LC-MS/MS)

Separation of digitonin-solubilized mitochondrial proteins on one-dimensional BN-PAGE and two-dimensional BN/SDS-PAGE was performed (Eubel et al., 2003), using a 4.5%-16% gradient. In-gel staining of Complex I or Complex IV activity was carried out as outlined previously (Sabar et al., 2005).

Gel spots were excised from 2D-BN/SDS-PAGE and sliced into smaller cubes (∼1mm) with a razor blade where necessary. Peptides in each gel piece were digested overnight with trypsin at 37°C and then extracted as previously described (Nelson et al., 2014). Following resuspension of digested peptides in 20 μL of 2% (v/v) acetonitrile and 0.1% (v/v) formic acid, samples were filtered before they were loaded onto a C18 high-capacity nano LC chip (Agilent Technologies) and eluted into Agilent 6550 Q-TOF with a 1200 series capillary pump as described previously (Nelson et al., 2014).

Results were searched against an in-house Arabidopsis database comprising ATH1.pep (release 10) from The Arabidopsis Information Resource (TAIR) and the Arabidopsis mitochondrial and plastid protein sets (the combined database contained a total of 33,621 protein sequences with 13,487,170 residues) using the Mascot search engine version 2.3. Search parameters include: error tolerances of 20 ppm for MS and 0.5 D for MS/MS, “max missed cleavages” set to 1, and variable modifications of oxidation (Met) and carbamidomethyl (Cys). Results were filtered with an ion cut-off score of 0.

### Analysis of mitochondrial peptides by liquid chromatography selective reaction monitoring mass spectrometry (LC-SRM-MS)

One hundred micrograms of mitochondrial protein was precipitated in 100% acetone overnight at −20°C and the pellets were washed with ice cold acetone for three times. Samples were alkylated, trypsin digested, desalted and cleaned as previously described (Petereit et al., 2020). The digested peptides were loaded onto an AdvanceBio Peptide Map column (2.1 mm × 250 mm, 2.7 μm particle size; part number 651750-902, Agilent), using a Thermo UltiMate 3000 RSLCnano System coupled to an Thermo Altis Triple Quadrupole MS. The column was heated to 55 °C, and the column flow rate was 0.4 ml/min. The binary elution gradient for HPLC was described previously (Le et al., 2022). The list of peptide transitions used for SRM–MS are provided in Table S2 with peak area data. Peak area of targeted peptides was determined using the Skyline software package version 21.2.0.425. Peptide abundances from each sample were normalized against voltage-dependent anion channel (VDAC).

### Analyses of organic acids and amino acids by LC-SRM-MS

Leaf discs or roots (∼25 mg) were collected at specified time points and immediately snap-frozen in liquid nitrogen. Metabolites were extracted as previously specified (Lee et al., 2021).

For LC-MS analysis of organic acids, sample derivatization was carried out based on a previously published method with modifications (Han et al., 2013). Samples were analyzed by an Agilent 1100 HPLC system coupled to an Agilent 6430 Triple Quadrupole (QQQ) mass spectrometer equipped with an electrospray ion source as described previously (Lee et al., 2021).

For amino acid quantification, dried samples were resuspended in 50 ml water as described previously (Le et al., 2021). Briefly, chromatographic separation was performed using Agilent Poroshell 120 HILIC-Z column, using mobile phases of 20 mM ammonium formate in water (solvent A) and 20 mM ammonium formate in acetonitrile (Solvent B). The elution gradient was 100% B at 0 min, 92% B at 1 min, 70% B at 10 min, 30% B at 10.5 min, 30% B at 12.5 min, 100% B at 12.5 min and 100% B at 25 min. The column flow rate was 0.4 mL/min; the column temperature was 35 °C, and the autosampler was kept at 10°C. The Agilent 6430 QQQ-MS was operated in positive ion mode in SRM mode. Data acquisition and LC-MS control were done using the Agilent MassHunter Data Acquisition software (version B06.00 Build 6.0.6025.4). The autosampler was kept at 10°C. The QQQ-MS was operated in SRM mode using the following operation settings: capillary voltage, 4000V; drying N2 gas and temperature, 11 L/min and 125 °C respectively; Nebulizer, 15 psi. Data analysis was carried out using MassHunter Quantitative Analysis Software (version 10.1, Build 10.1.733.0). Metabolites were quantified by comparing the integrated peak area with a calibration curve obtained using authentic standards, and normalised against fresh weight and internal standards.

### Analyses of nucleotides by LC-SRM-MS

Absolute quantitation of ADP and ATP by LC-MS was carried out according to previous reports (Straube et al., 2021) with slight modifications. Briefly, approximately 25-mg roots or 50-mg leaf discs were collected and immediately snap-frozen in liquid nitrogen. Samples were ground to a fine powder and 1 mL of ice-cold 15% TCA solution supplemented with ^13^C5,15N5-AMP as an internal standard. Following centrifugation at 24,000 x *g* for 10 min (4°C), 1 mL 78/22 DCM/TOA was added to the supernatant. The mixture was then vortexed and centrifuged at 5,000 x g for 2 min. The upper phase was collected and diluted in 1 mL H_2_O and 5 µL 0.5% acetic acid. The resulting mixture was applied to a Strata X-AW SPE cartridge (pre-equilibrated with 1 mL methanol, 1 mL 2/25/73 formic acid/methanol/H_2_O, and 1 mL 10 mM ammonium acetate pH 4.5) and the flow-through was discarded. The cartridge was then washed with 1 mL 1 mM ammonium acetate (pH 4.5) and 1 mL methanol before nucleotides were eluted with 0.5 mL 20/80 ammonia/methanol twice. The eluate was transferred to a new tube and dried using a SpeedVac.

Dried samples were resuspended in 100 µL water. Chromatographic separation was performed using Agilent Poroshell 120 HILIC-Z column, using mobile phases of 5 mM ammonium acetate pH 9.0 (solvent A) /5 mM ammonium acetate, 90% ACN (Solvent B). Solvents A and B were supplied with 0.1% (v/v) Infinity Lab deactivator additive (Agilent) to improve peak shapes. The elution gradient was 100% B at 0 min, 60% B at 5 min, 30% B at 5.5 min, 30% B at 7 min, 15% B at 8.5 min, 100% B at 9 min and 100% B at 22 min. The column flow rate was 0.3 mL/min; the column temperature was 35 °C, and the autosampler was kept at 10°C. Data acquisition and LC-MS control were carried out using the Agilent MassHunter Data Acquisition software (version B06.00 Build 6.0.6025.4). The autosampler was kept at 10°C. The QQQ-MS was operated in SRM mode in positive ion polarity using the following settings: capillary voltage, 4000V; drying N2 gas and temperature, 11 L/min and 125 °C respectively; Nebulizer, 15 psi. All optimised SRM transitions for each target were listed in Table S4. Data analysis was carried out using MassHunter Quantitative Analysis Software (version 10.1, Build 10.1.733.0). Metabolites were quantified by comparing the integrated peak area with a calibration curve obtained using authentic standards, and normalised against fresh weight and internal standards.

### Statistical Analysis

All statistical analyses were performed with RStudio (including Student’s t-test, ANOVA with Tukey Posthoc analysis, Shapiro-Wilk normality test and Levene’s Test). Statistical tests and replicate numbers are as indicated in figure legends.

## Author contributions

CPL performed most of the mitochondrial experiments and LC-MS analysis of organic acids, amino acids and nucleotides; XHL performed mitochondrial proteomics, CIV assays, membrane potential measurements and whole plant FCCP sensitivity assays; RG performed phylogenetic studies, sequences searches, HMM analysis and interpretations; JAC and SJR screened *enod93* mutants, generated complementation and overexpression lines and performed preliminary phenotype assessments; AHM designed experimental strategies and wrote the manuscript with CPL. All authors contributed to the revision of the manuscript and discussion.

## Acknowledgements

This work was supported by funding from the Australian Research Council (CE140100008; FL200100057) to AHM and by the Natural Sciences and Engineering Research Council of Canada (NSERC) to SJR. We thank Prof Hans-Peter Braun, Dr Jennifer Senkler, Dr Holger Eubel and Dr Nils Rugen from the University of Hannover for discussions on the protein data and their complexome maps of Arabidopsis mitochondrial proteins (complexomemap.de). Peptide quantitation in this work was performed as a service by Dr Elke Ströher from the WA Proteomics Facility as a node of Proteomics Australia, supported by infrastructure funding from the Western Australian State Government in partnership with Bioplatforms Australia under the Commonwealth Government National Collaborative Research Infrastructure Strategy.

## Supplemental Tables

**Table S1 –** Peptide mass spectrometry identifications of ENOD94 in gel spots for BN-PAGE

**Table S2 –** Peptide MRMs quantification for different mitochondrial components

**Table S3 –** PCR Primers used in the study

**Table S4 –** SRMs for nucleotides used in the study

## Supplemental Figure Legends

**Figure S1.**
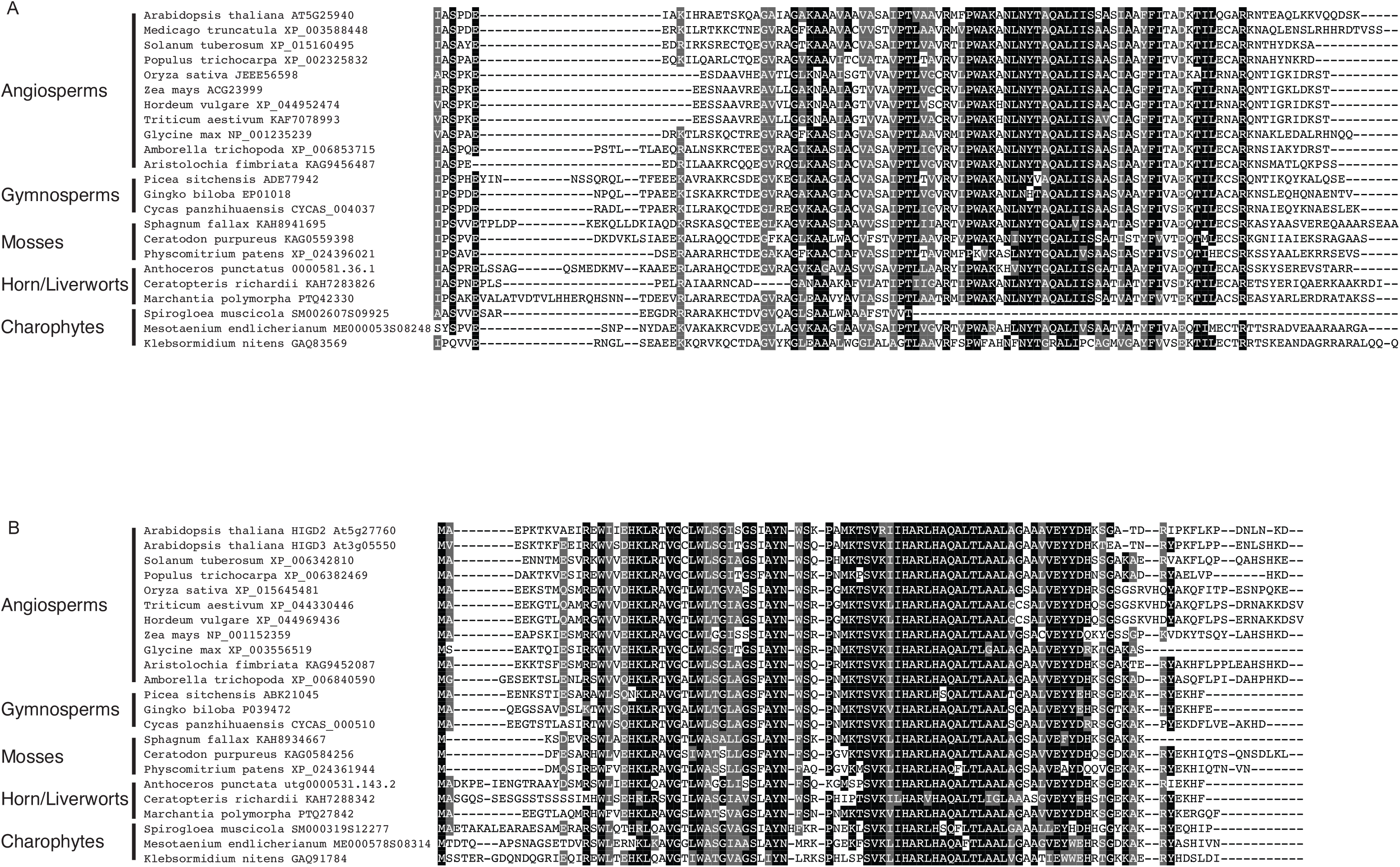
ENDO93 and Rcf2 homologs are widespread across plants. A) Multiple alignment of ENOD93 homologs from diverse angiosperms, gymnosperms, mosses, hornworts, liverworts, and charophyte algae. A small and poorly aligned N-terminal portion was removed for presentation. B) Multiple alignment of Rcf2 homologs from diverse angiosperms, gymnosperms, mosses, hornworts, liverworts, and charophyte algae. Shaded characters indicate positions with ≥80% sequence identity (black) or similarity (gray), defined according to amino acids belonging to groups with similar physicochemical properties (GAVLI, FYW, CM, ST, KRH, DENQ, P).

**Figure S2.**
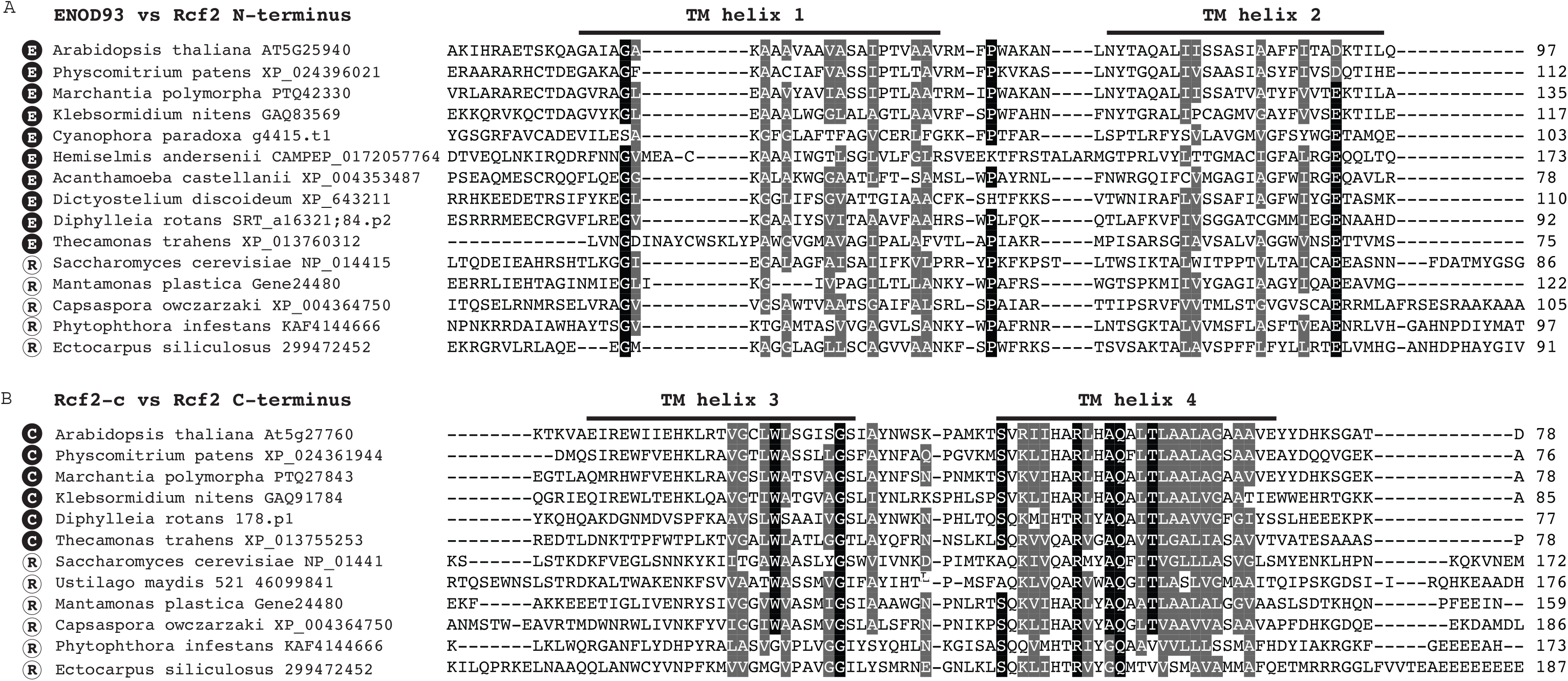
Phylogenetically broad protein alignments demonstrate homology between ENOD93 and Hig1-domain proteins with the N- and C-terminus of Rcf2, respectively. A) Partial multiple alignment of putative ENOD93 homologs with the N-terminal region of putative Rcf2 from diverse eukaryotes. B) Partial multiple alignment of Hig1-domain proteins with the C-terminal region of putative Rcf2 homologs from diverse eukaryotes. Transmembrane helices of yeast Rcf2 are marked with solid lines according to Zhou et al. (2021). Shaded characters indicate positions with ≥80% sequence identity (black) or similarity (gray), defined according to amino acids belonging to groups with similar physicochemical properties (GAVLI, FYW, CM, ST, KRH, DENQ, P). Dark circles with an ‘E’ indicated putative Enod93 homologs; dark circles with an ‘H’ indicated Hig1-domain proteins equivalent to the C-terminal region of Rcf2; and white circles with an ‘R’ indicate full-length Rcf2 homologs.

**Figures S3.**
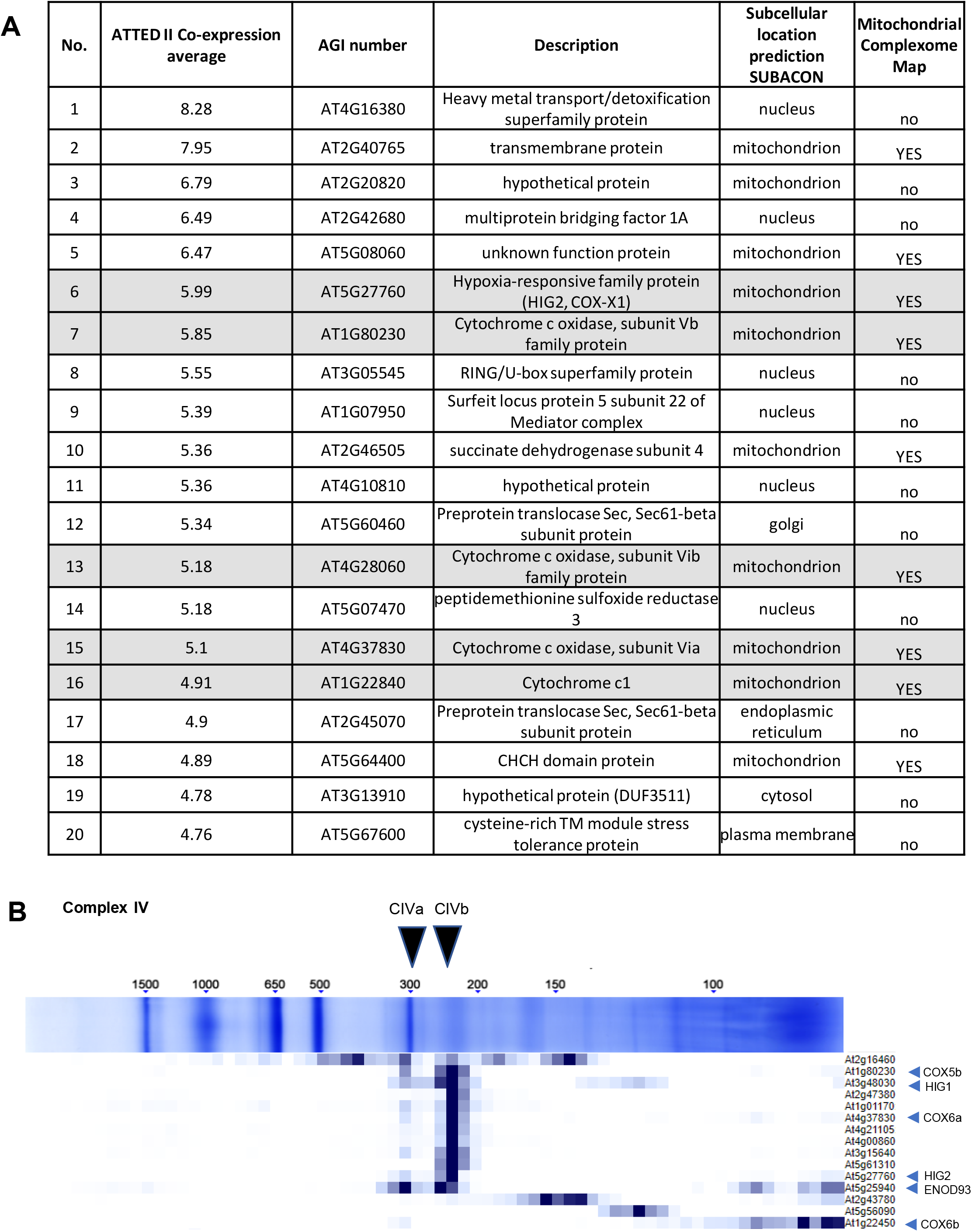
Co-expression and intra-mitochondrial localisation of ENOD93. (A) The 20 genes most co-expressed with *ENOD93* in Arabidopsis from the co-expression function in ATTED-II (Obayashi et al., 2022), the predicted subcellular localization of the encoded proteins as determined by SUBAcon (Hooper et al., 2014) and their presence/absence in the Arabidopsis mitochondria complexome map (Senkler et al., 2017). Genes for subunits of cytochrome c oxidase are highlighted. (B) Reproduction of the clustering of the relative abundance of ENOD93 peptides with those of other complex IV subunits in the Arabidopsis mitochondria complexome map (Senkler et al., 2017) using complexomemap.de, and annotation of the size of the two intact versions of CIV; CIVa and CIVb (Millar et al., 2004), and the location of ENOD93, HIG1, HIG2, COX5b, COX6b and COX6a.

**Figure S4.**
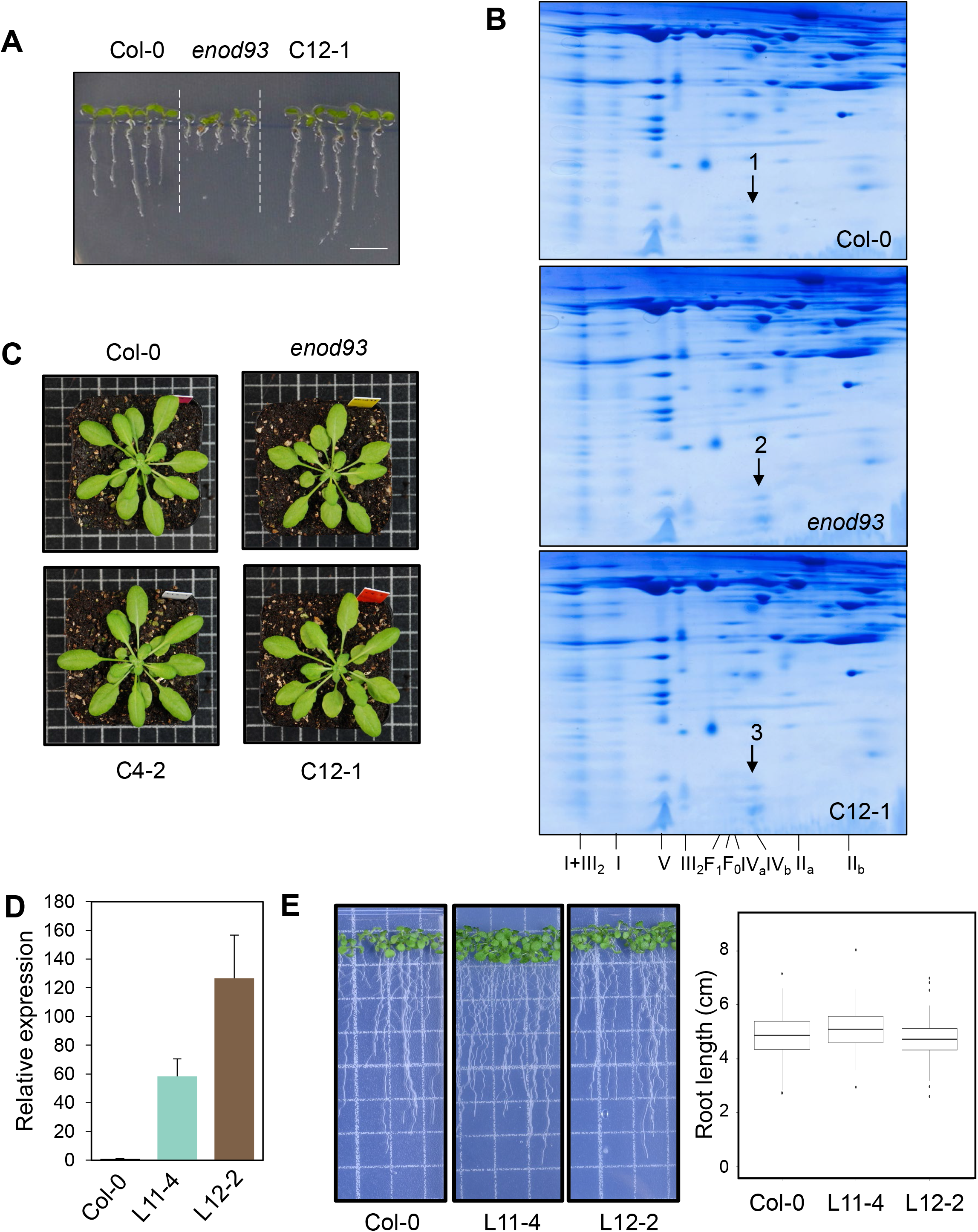
Phenotype of *enod93*, complemented and overexpression lines and identification of ENOD93. (A) Representative image of vertically grown 8-day-old seedlings of Col-0, *enod93* and complemented lines on an agar plate under long day condition. (B) 2D-blue-native/SDS-PAGE separation of mitochondrial proteins from the three genotypes. Roman numerals correspond to the locations of respiratory complexes. Gels were visualised by Coomassie Blue. Location of protein spots containing ENOD93 marked by arrows were verified by LC-MS/MS, with the corresponding numbers correlate with Table S1. (C) Phenotype of Col-0, *enod93*, C4-2 and C12-2 lines in ∼4-5 weeks of long-day light growth conditions. (D) Expression levels of Col-0 and ENOD93 overexpression lines (L11-4 and L12-2) as determined by qPCR (n = 3). (E) Representative image of vertically grown seven-day-old seedlings of Col-0 and ENOD93 overexpression lines on an agar plate under long day condition (left). Root length of seedlings is shown on the right. No significant change in root length was found based on Kruskal-Wallis rank sum test

**Figure S5.**
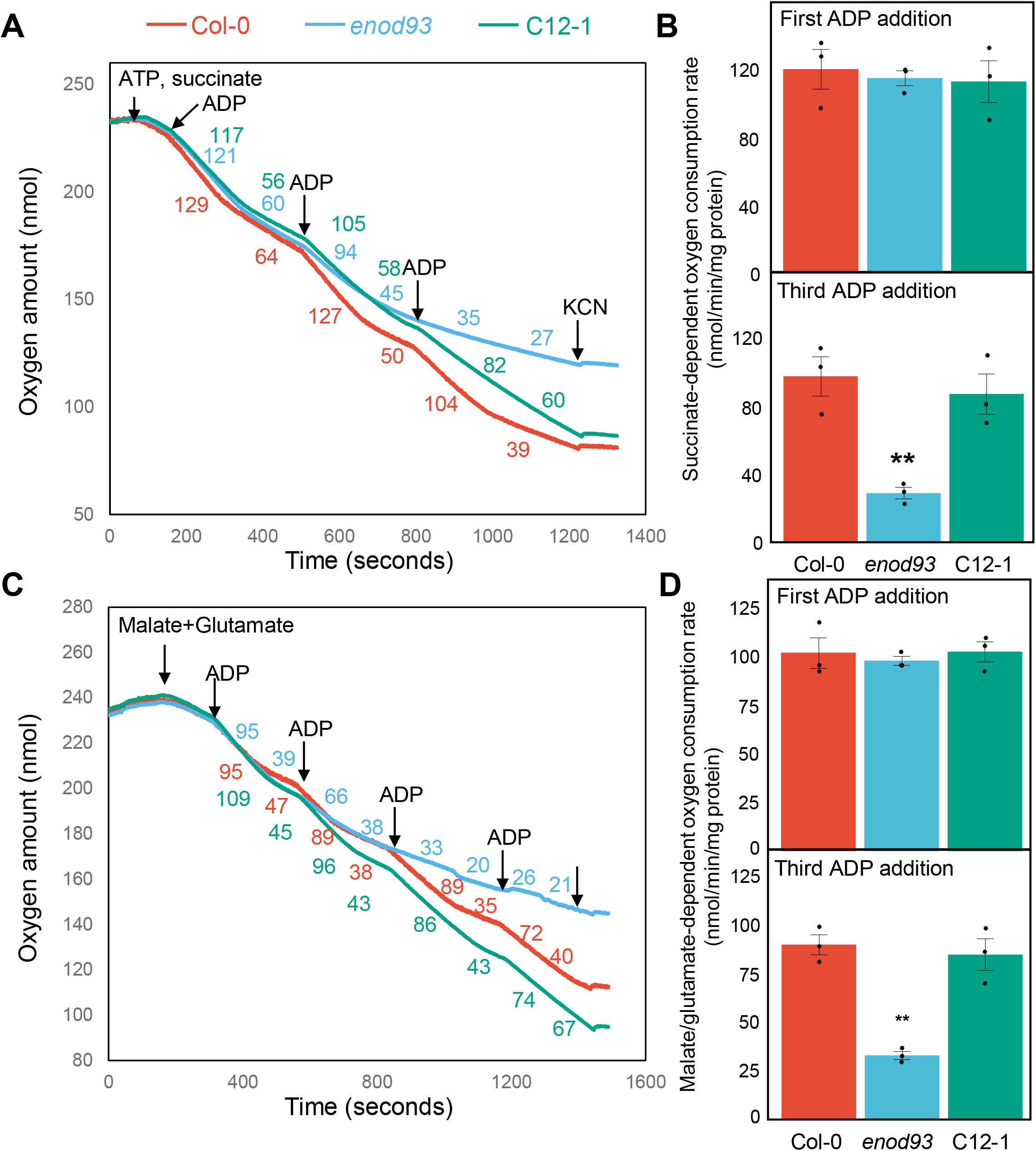
Succinate- or malate/glutamate-dependent respiration by isolated *enod93* mitochondria. (A) Representative trace illustrating succinate-stimulated oxygen consumption by mitochondria purified from Col-0, *enod93* and C12-1 (a complementation line) seedlings. Oxygen consumption rates in response to the successive treatments are shown in coloured numerical values. Substrate additions to all samples are indicated by arrows with the following concentrations: 5 mM succinate, 0.5 mM ATP, 0.1 mM ADP and 1 mM KCN. (B) State III oxygen consumption rates for purified mitochondria energized with succinate in response to the first (upper panel) and third (lower panel) ADP addition as indicated in (A). (C) Representative trace illustrating malate/glutamate-stimulated oxygen consumption by mitochondria purified from Col-0, *enod93* and C12-1 seedlings. Oxygen consumption rates in response to the successive treatments are shown in coloured numerical values. Substrate additions to all samples are indicated by arrows with the following concentrations: 10 mM malate, 10 mM glutamate, 0.1 mM ADP and 1 mM KCN. (D) State III oxygen consumption rates for purified mitochondria energized with malate and glutamate in response to the first (upper panel) and third (lower panel) ADP addition as indicated in (C). (E) Oxygen consumption rates for purified mitochondria energized with succinate in response to ADP addition with and without 0.3 mM malonate. (F) Percentage of safranin uptake as a measure of membrane potential in mitochondria from WT and *enod93* after multiple ADP additions during malate/pyruvate-stimulated respiration in the presence of 3 mM ferricyanide as an alternative oxygen acceptor to bypass cytochrome c oxidase. Data represents mean ± S.E. with overlaid individual data points as dots (n = 3). Asterisks indicate a significant change as determined by one-way ANOVA with Tukey post-hoc test (* p < 0.05; ** p < 0.01).

**Figure S6.**
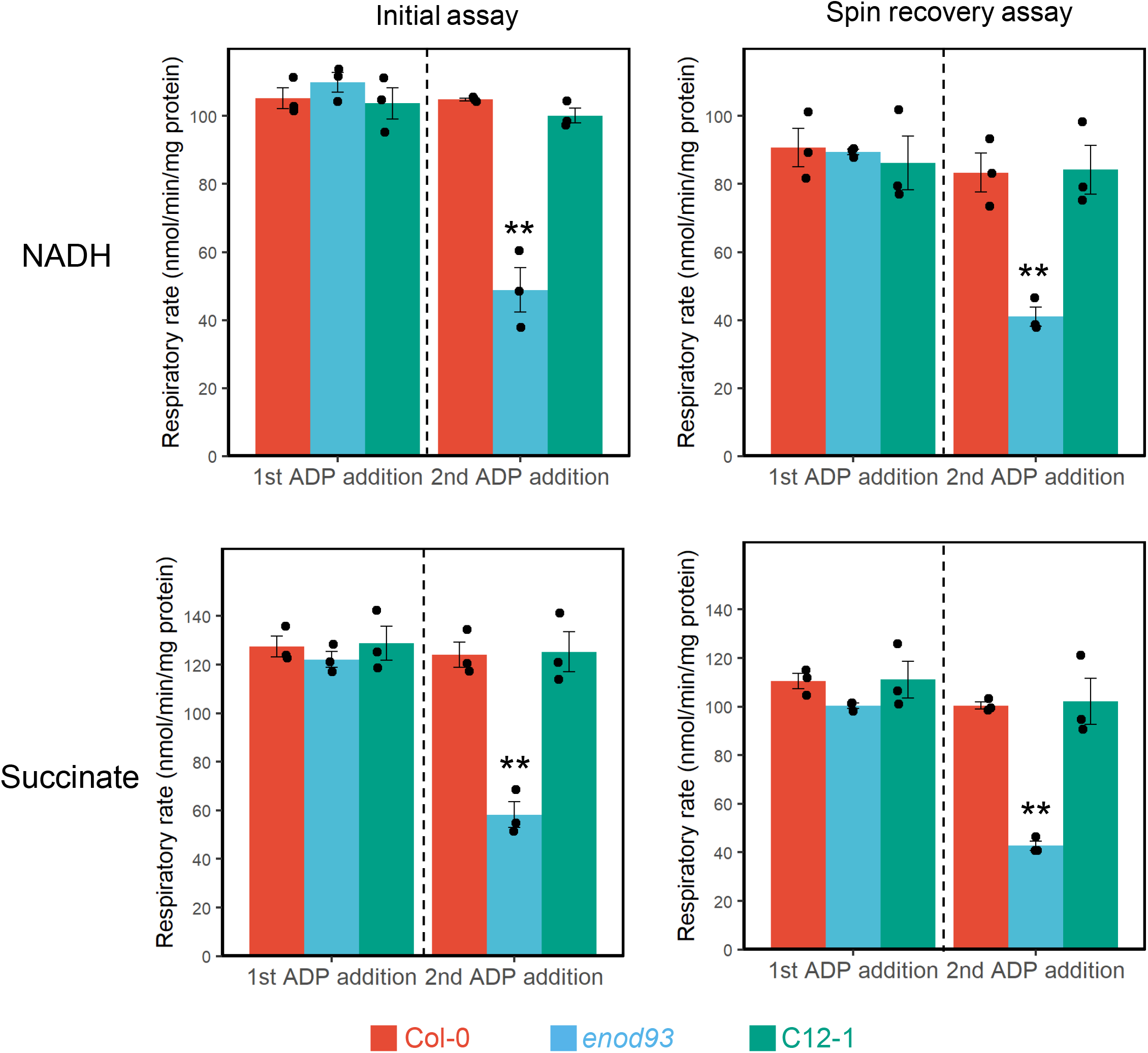
Spin recovery assays of respiratory rate in isolated *enod93* mitochondria. For Initial assay, NADH- (upper panels) or succinate- (lower panels) dependent state 3 respiration measurements were carried out on freshly purified mitochondria. Mitochondrial fractions were collected and subjected to a few rounds of washes with respiration medium without substrates to dilute or remove substrates, products and ATP. Respiration measurements were then repeated on washed mitochondria fractions, as shown under the spin recovery assay heading. Data represents mean ± S.E. with overlaid individual data points as dots (n = 3). Asterisks indicate a significant change in Col-0 vs *enod93* and *enod93* vs C12-1 comparisons as determined by one-way ANOVA with Tukey post-hoc test (* p < 0.05; ** p < 0.01).

**Figure S7.**
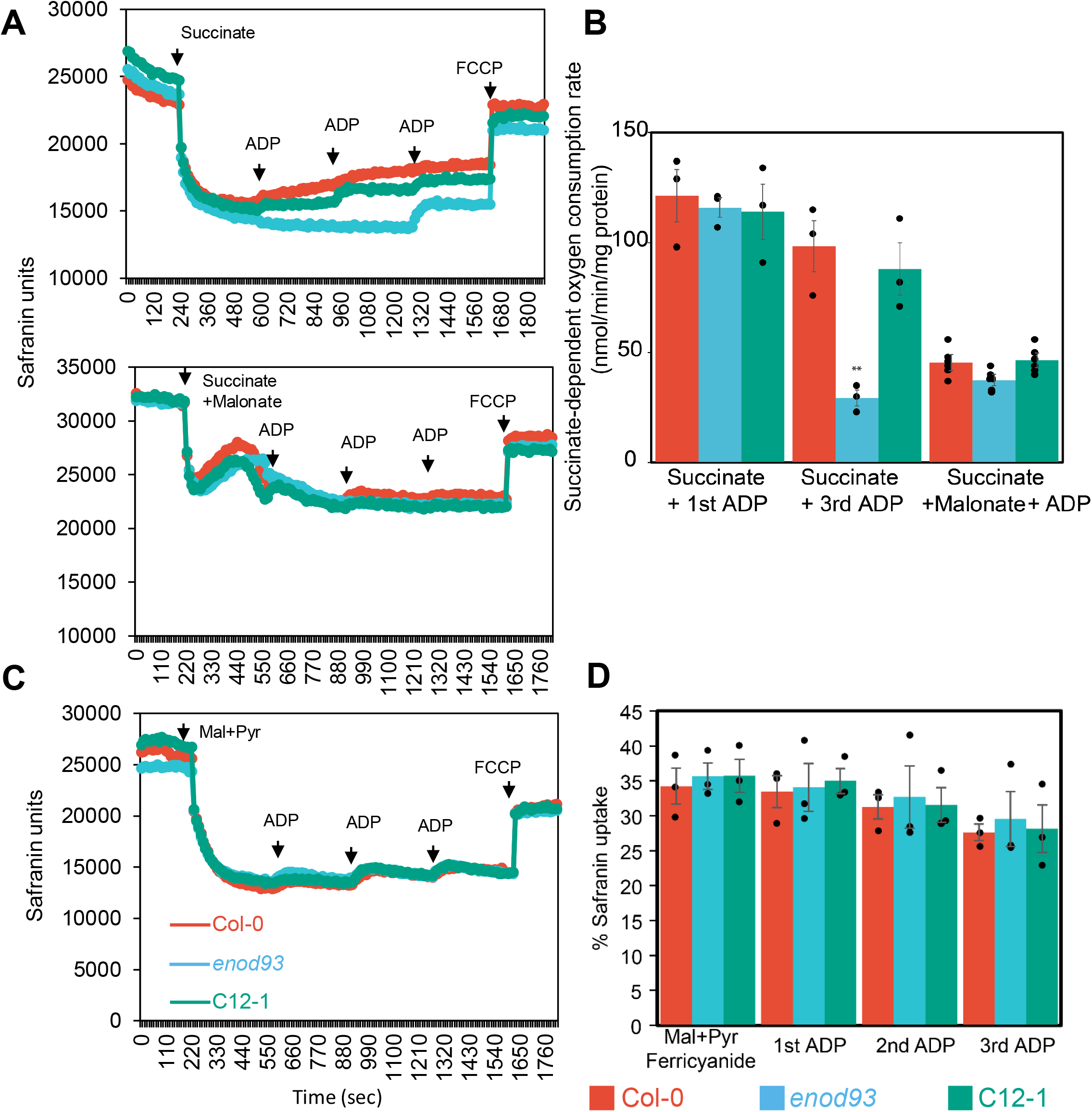
Effect of lowering membrane potential and complex IV dependence based on differences between WT, *enod93* and complemented line mitochondrial bioenergetics. (A) Representative trace of safranin uptake as a measure of membrane potential in mitochondria from Col-0, enod93 and C12-1 after multiple ADP additions during succinate-stimulated respiration with/without 0.3mM malonate as a complex II inhibitor. (B) Oxygen consumption rates for purified mitochondria energized with succinate in response to ADP addition with and without 0.3 mM malonate. Asterisks indicate a significant change as determined by Student’s t-test (** p < 0.01). (C) Representative trace of safranin uptake as a measure of membrane potential in mitochondria from Col-0, enod93 and C12-1 after multiple ADP additions during malate/pyruvate-stimulated respiration in the presence of 3 mM ferricyanide as an alternative oxygen acceptor to bypass cytochrome c oxidase. (D) % safranin uptake in repeated experiments with data representing mean ± S.E. with overlaid individual data points as dots (n = 3). No significant difference was determined by Student’s t-test.

**Figure S8.**
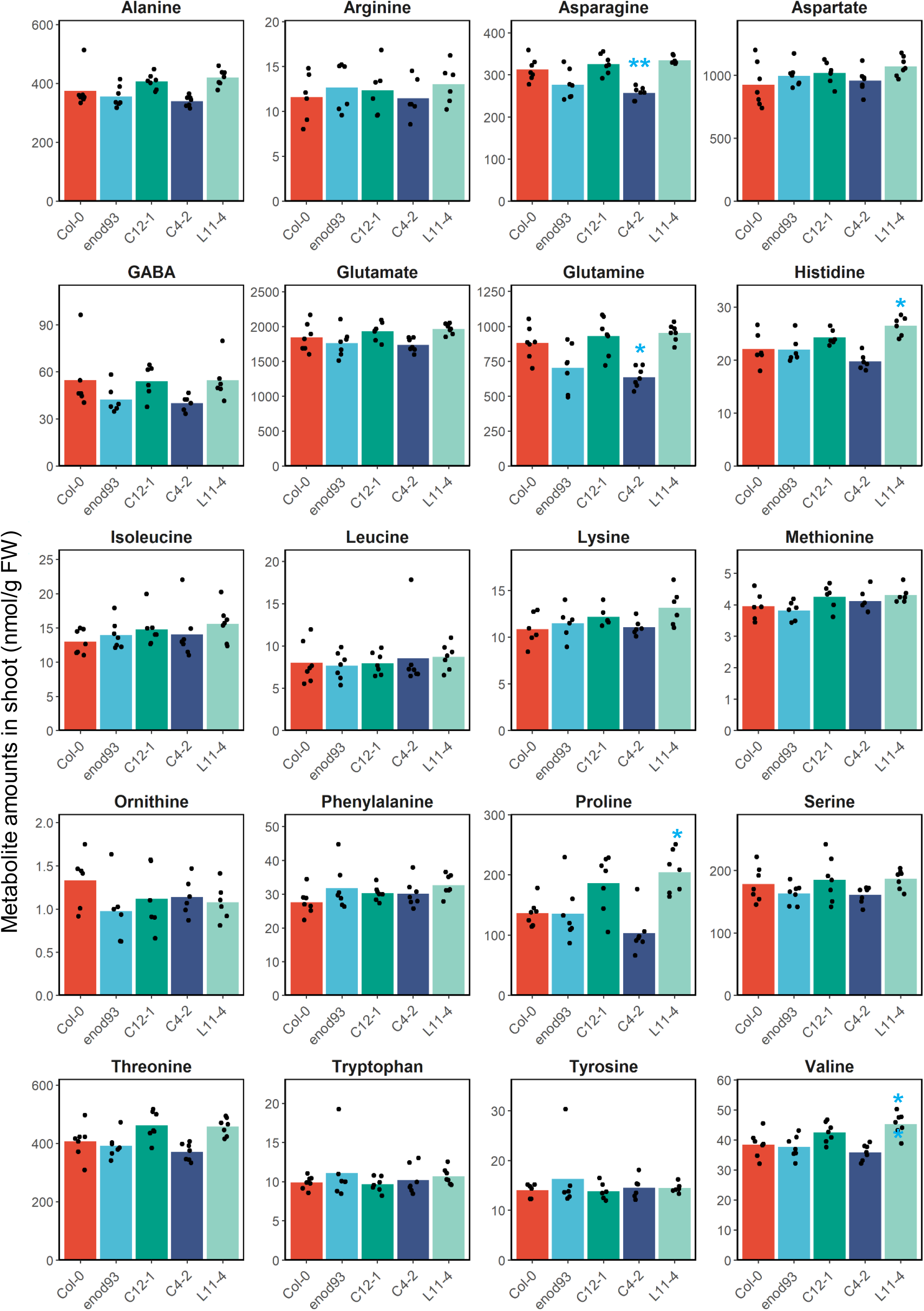
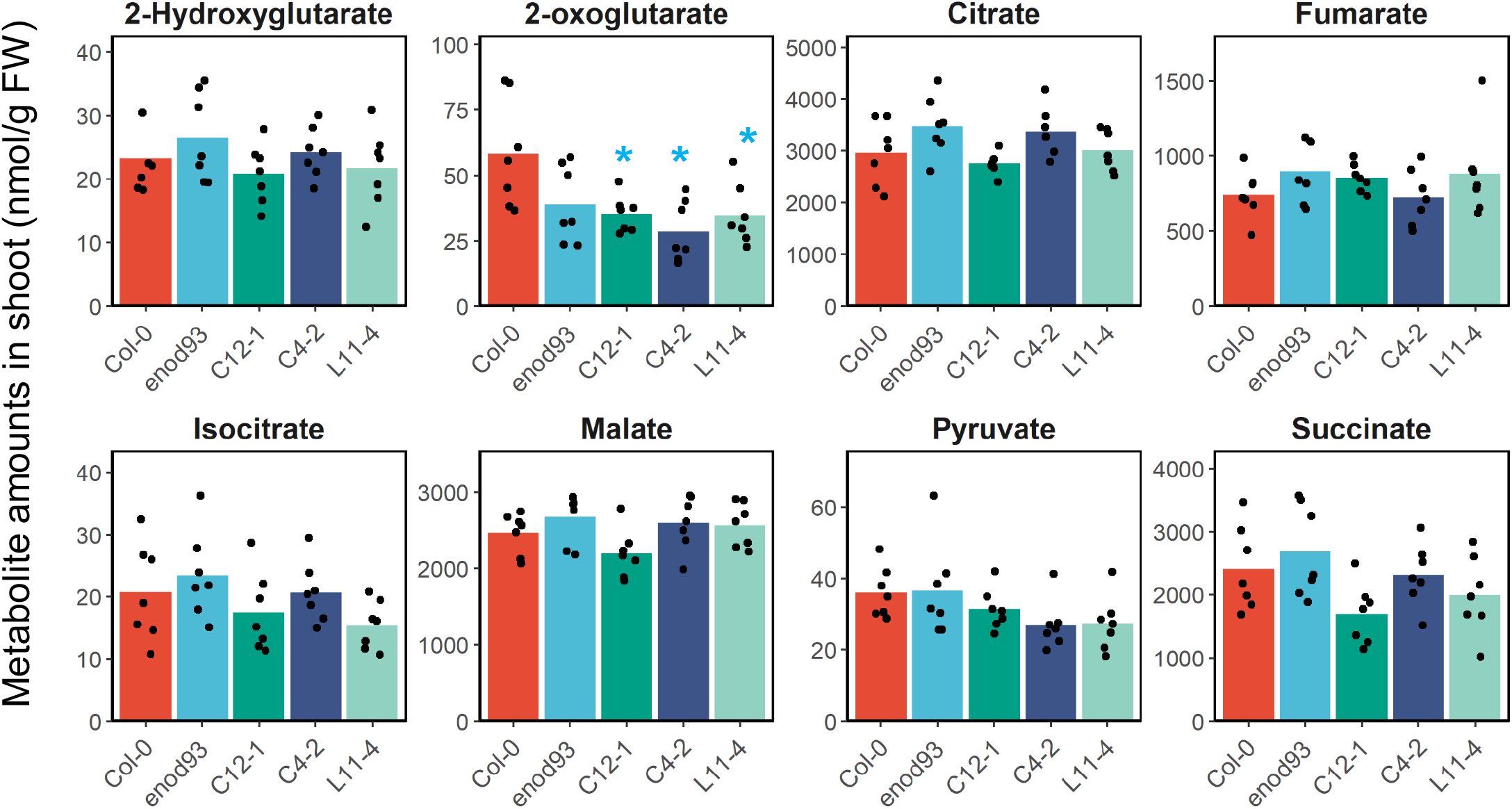
Leaf organic and amino acid content in WT, *enod93,* complemented and over-expression genotypes. Plants were grown under long day conditions for 5 weeks and leaf discs were collected 4 hours after dark shift. Metabolites in this figure were analysed by LC-MS. Data shown are mean with overlaid individual data points as dots (n ≥ 6) in nmol per gram fresh weight (FW), and asterisks indicate a significant change relative to Col-0 as determined by Student’s t-test (** p < 0.01, * p < 0.05). *enod93* complemented lines C12-1 and C4-2 are shown. Overexpressed *ENOD93* line L11-4 is shown.

**Figure S9.**
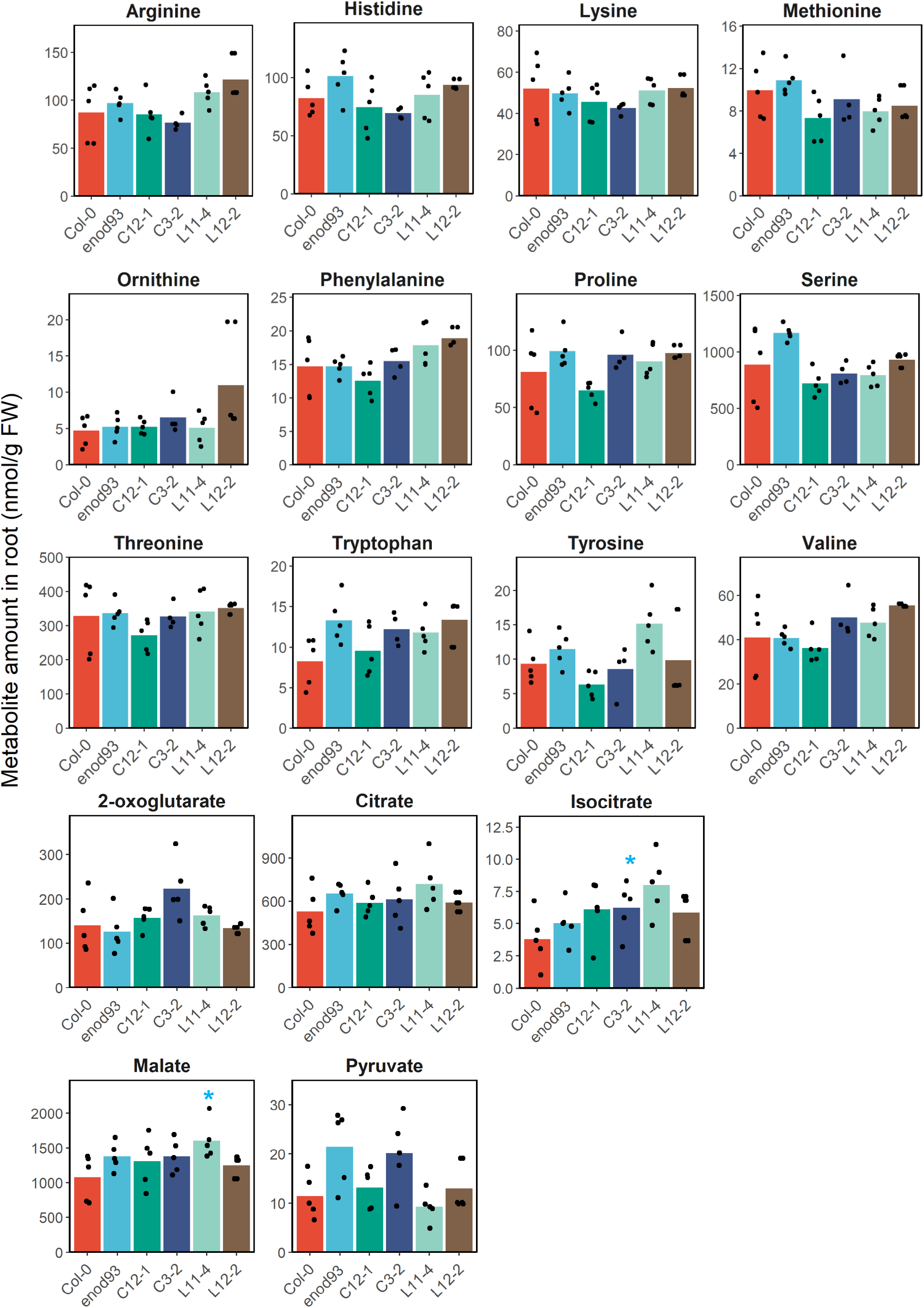
Root organic and amino acid content in WT, *enod93*, complemented and over- expression genotypes. Roots from a pool of seven-day-old seedlings were collected in liquid nitrogen and extracted metabolites were analysed by LC-MS. Data shown are mean with overlaid individual data points as dots (n ≥ 5) in nmol per gram fresh weight (FW), and asterisks indicate a significant change relative to Col-0 as determined by Student’s t-test (* p < 0.05). *enod93* complemented lines C12-1 and C3-2 and overexpressed *ENOD93* lines L11-4 and L12-2 are shown. Averages shown in bar graphs, n=4, *p<0.05, **p<0.01. Showing organic and amino acid contents not shown in Figure 8.

**Figure S10.**
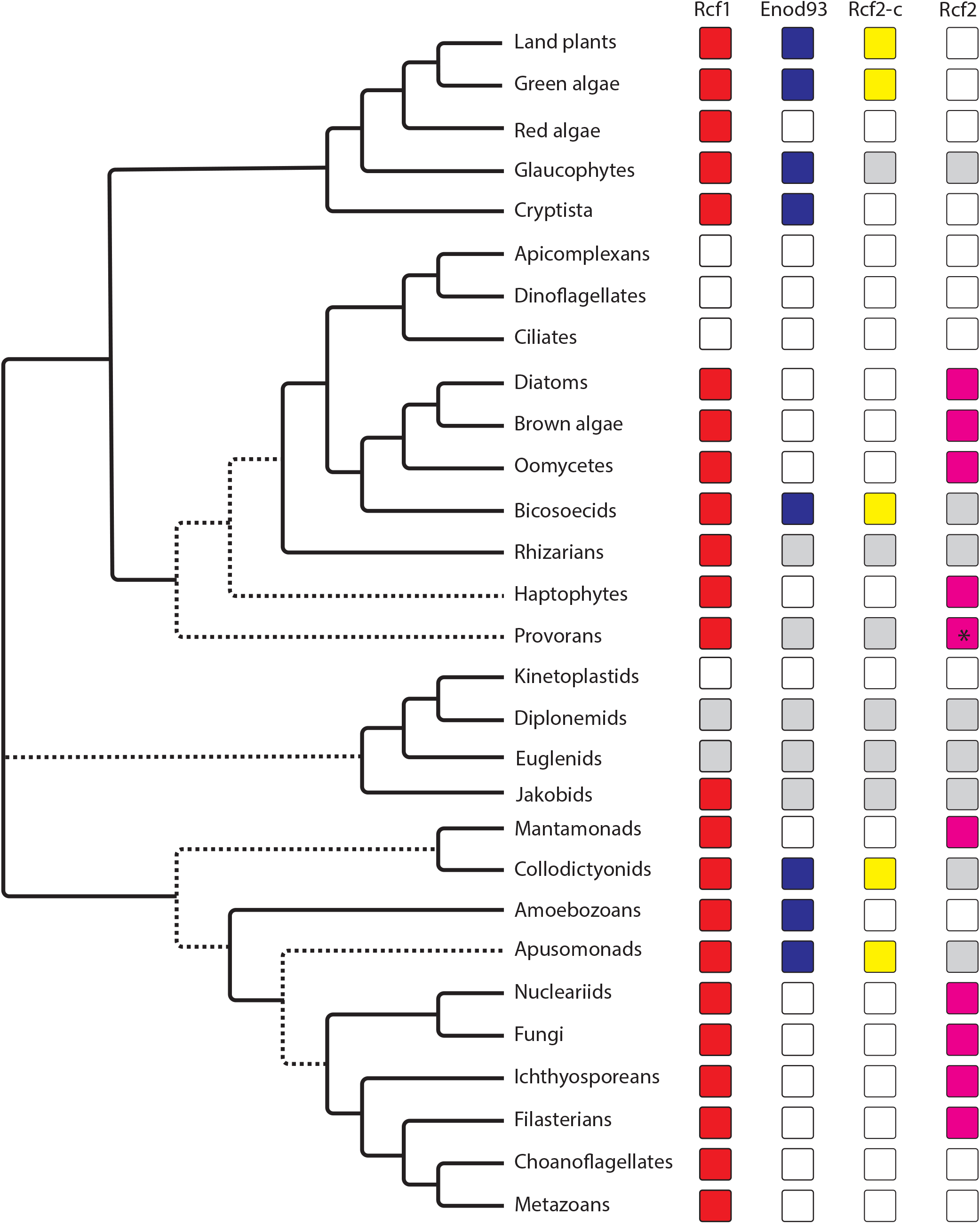
Complex distribution of plant- and fungal-type Rcf2 homologs across the breadth of eukaryotes. The distribution of plant- and fungal-type Rcf2 does not precisely mirror eukaryotic phylogeny, suggesting the multiple gene fusions and/or fissions have occurred. Presence of Rcf1 (red), plant-type Rcf2 – consisting of Enod93 (blue) and Rcf2-c (yellow) – and fungal-type (magenta) Rcf2 are shown. Empty boxes indicate an inability to find homologs in listed groups. Boxes shaded light gray also indicate that homologs were not found, but that insufficient genomic resources are available to be confident of absence. Dashed lines denote tentative associations between major eukaryotic lineages. An asterisk is shown for fungal-type Rcf2 in Provorans because protein domains are fused, but in the opposite orientation of fungal Rcf2.

